# Understanding DNA interactions in crowded environments with a coarse-grained model

**DOI:** 10.1101/2020.06.08.140434

**Authors:** Fan Hong, John S. Schreck, Petr Šulc

## Abstract

Nucleic acid interactions under crowded environments are of great importance for biological processes and nanotechnology. However, the kinetics and thermodynamics of nucleic acid interactions in a crowded environment remain poorly understood. We use a coarse-grained model of DNA to study the kinetics and thermodynamics of DNA duplex and hairpin formation in crowded environments. We find that crowders can increase the melting temperature of both an 8-mer DNA duplex and a hairpin with a stem of 6-nt depending on the excluded volume fraction of crowders in solution and the crowder size. The crowding induced stability originates from the entropic effect caused by the crowding particles in the system. Additionally, we study the hybridization kinetics of DNA duplex formation and the formation of hairpin stems, finding that the reaction rate *k*_on_ is increased by the crowding effect, while *k*_off_ is changed only moderately. The increase in *k*_on_ mostly comes from increasing the probability of reaching a transition state with one base pair formed. A DNA strand displacement reaction in a crowded environment is also studied with the model and we find that rate of toehold association is increased, with possible applications to speeding up strand displacement cascades in nucleic acid nanotechnology.

## I. INTRODUCTION

The interactions between DNA/RNA strands are controlled by orthogonal base pairing of adenine (A) to thymine (T) and cytosine (C) to guanine (G), and are essential for fundamental cellular activities and practical molecular therapeutic and diagnostic purposes, such as gene replication^1^, gene regulation^2^ and diagnostics^3^ as well as anti-sense oligonucleotide drugs^4^. Furthermore, recent years have also witnessed the emerging field of DNA nanotechnology, which uses DNA to build complex designed molecular nanostructures and molecular machines by taking advantage of its programmability and predictable interactions^5–8^, allowing for unprecedented precise control of structure and dynamic behavior at the nanoscale. The properties of DNA strand interactions, such as the thermodynamics and kinetics of duplex formation, determine the assembly efficiency and stability of the DNA structures that affect cellular functions, anti-sense drug efficiency, and the performance of designed molecular machines^1–4,9^. Consequently, a quantitative understanding of the biophysical properties of DNA interactions is required to rationalize fundamental cellular functions, as well as to improve nucleic acid-based biotechnologies.

Biophysical properties of DNA and RNA have been well-studied in diluted aqueous solutions^10,11^, and have been successfully applied to DNA (and RNA) secondary structure prediction^10,12,13^ and molecular probe design for molecular diagnostics and next-generation sequencing^14,15^. However, the interaction parameters extracted from a diluted solution do not necessarily correspond to DNA and RNA interactions *in vivo*. In living cells, the environment is occupied with macro-molecules, cell organelles, and relatively small molecules, such as metabolites and osmolytes, occupying 10% - 40% of the total volume^16–18^, resulting in highly crowded and complicated conditions^19,20^. Researchers have found that RNA and DNA have significant differences in the thermodynamic and folding properties between the crowded environments and aqueous solutions^21–24^. This macromolecular crowding in a cellular environment has a significant influence on both the intramolecular and inter-molecular interactions by changing the energetic and transport properties of the molecules in the crowding environment.

To improve the performance of nucleic acid-based technologies and our understanding of DNA/RNA-related cellular activities, comprehensive studies of DNA interactions in the crowded environment are necessary and of great importance. Generally, the kinetic and thermodynamic properties of DNA/RNA are investigated through single-molecule and UV melting experiments under crowded conditions achieved in *in vitro* by introducing crowding agents (such as PEG, sucrose, urea, dextran and others^25^) into the solution. The comparison of the results between the aqueous solution and the crowded environment is then used to infer principles governing the influence of the crowders on the biophysical properties.

DNA-DNA interactions are complicated processes involving diffusion, nucleation, and zippering steps^26^. Although experimental studies can capture the overall results, which can be used to create a simplified physical model to describe the crowding effects^23,25,27–29^, the individual steps of the hybridization or duplex formation cannot be easily captured by the experimental methods. As a complement to the experiments, computational modeling, such as molecular dynamics (MD), can provide more detailed information about the process of DNA hybridization in a crowded environment. Ideally, a fully atomistic MD simulation would be used to simulate the hybridization process, but the time-scales involved in such processes are limited by rare events (such as the creation and breaking of bonds) make such a study infeasible. To address these challenges, a coarse-grained model is required to reduce the computational complexity while at the same time retaining the key information about the molecule. Prior computational studies have investigated the entropic stabilization of a folded RNA state in a crowded environment^30,31^, and the role that crowders play in polymer looping kinetics^20^.

Here we use a coarse-grained DNA model to study the DNA interactions under various crowding conditions. The oxDNA model^32–34^ is a coarsegrained DNA model that captures the structural, thermodynamic and mechanical properties of both single-stranded DNA and double-stranded DNA. Where available, the model has been able to quantitatively reproduce experimental measurements of DNA properties and has been successfully applied to the study of duplex and hairpin formation^26,32,35^, DNA behavior under a pulling force and torque^36,37^, DNA origami assembly^38–40^, properties of polyhedral and tile-based nanostructures^41–43^, and other complex processes such as strand displacement kinetics^44^ and DNA walkers^9,45^. We extend the oxDNA model by introducing inert crowding particles, which are represented by spheres interacting with excluded volume (schematically shown in Fig. 1). In this work, we focus purely on the entropic effect of the excluded volume interaction on the thermodynamics and kinetics of the interactions by neglecting other interactions between the crowders and DNA, such as electrostatic attraction or repulsion.

**FIG. 1.**
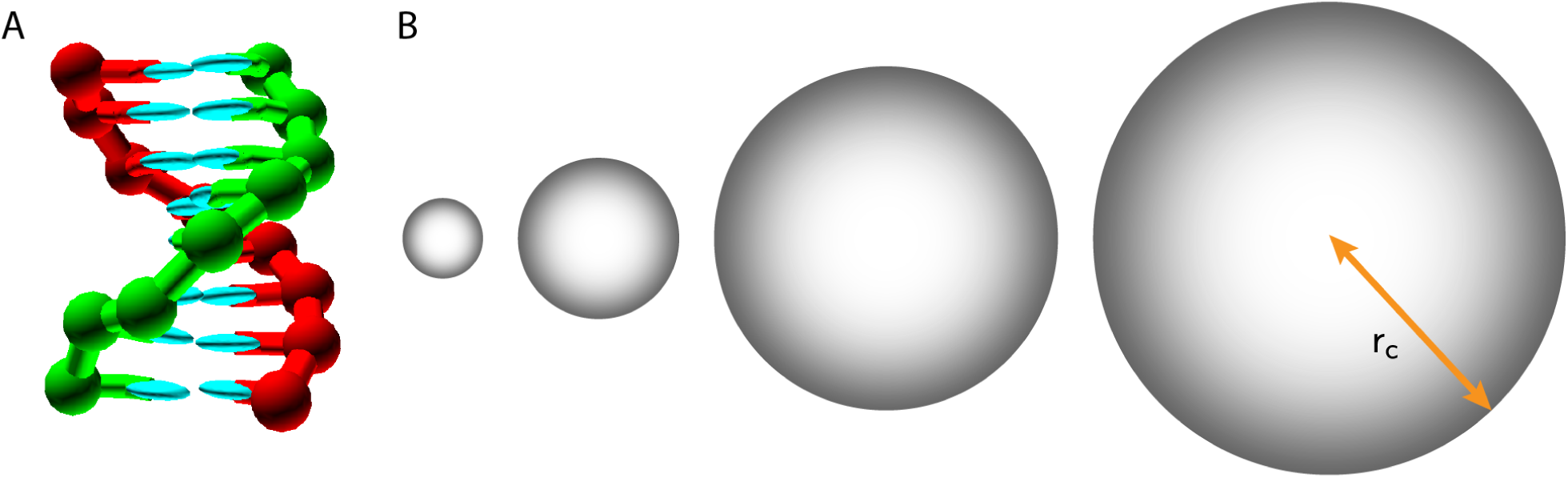
The crowder-oxDNA model. (A) A DNA 8-mer duplex configuration, as represented in the oxDNA model. (B) The crowder particles with different radii *r*c corresponding to the crowder sizes studied in the simulations, shown in relative size comparison with the DNA 8-mer.

With the extended coarse-grained model, we first study the thermodynamics of a DNA 8-mer duplex and hairpin with a stem of 6-nt and different loop lengths. We find that the crowders increase the melting temperature of both the DNA duplex and the hairpin. Increasing the volume fraction stabilizes the compact state (formed duplex or hairpin) with respect to the unbound state. The increased stability originates from the entropy change caused by the crowding particles in the system. For a fixed volume fraction, we observe smaller crowders having a much higher stabilizing effect than larger crowders, and is due to their higher number density. We fit an analytical formula based on scaled particle theory (SPT)^19,46–48^ to our thermodynamic data, and find semi-quantitative agreement in predicted change in binding free energy, with differences observed for smaller crowder sizes and higher volume occupied by crowders, where the approximations made by SPT are no longer applicable.

Additionally, we perform kinetic simulations of duplex and hairpin formation with individual nucleotide resolution. The rate-determining step of the two transitions is the nucleation step which involves the first hydrogen bond forming between the bases. We find that crowding effects generally favor the association rate *k*_on_, while the dissociation rate *k*_off_ is only weakly affected. The increase in *k*_on_ comes mostly from increasing the probability of reaching the transition state with one base pair formed, thus accelerating the association kinetics. Finally, we study a strand displacement reaction^44^ under different crowding conditions. We find that the binding to the toehold region is also increased in crowded environments. As a result, crowded environments could potentially be used to enhance strand displacement cascades rates in nanotechnological applications.

## II. METHODS

### A. Extending the oxDNA model to include spherical crowders

OxDNA is a coarse-grained model that treats a DNA strand as a chain of rigid bodies, with each rigid body representing a single nucleotide. The interaction potentials in the model, schematically illustrated in Fig. 1A and Fig. S1, have been parameterized to reproduce basic structural, thermodynamic, and mechanical properties of double-stranded and single-stranded DNA^32,34,49^. Each nucleotide has one interaction site to represent the backbone and two for the base (one for a stacking interaction and for a hydrogen-bonding interaction). Hydrogen-bonding and stacking interactions drive the formation of duplexes. The model has been shown to accurately represent the properties of duplexes as well as single-stranded DNA^26,32,49,50^.

We introduce here the extended crowder-oxDNA model. The crowders are modeled to mimic hard spheres, each having a given radius of *r*_*c*_, as seen in Fig. 1B. Crowder particles are assumed to interact with each other and with the nucleotides in the DNA strands through an excluded volume interaction only, as is described in detail in the Supporting Information. As the molecular dynamics simulations require the potentials to be differentiable, we use a strongly repulsive Lennard-Jones potential for when crowders come into contact with each other or with nucleotides, and no interaction if they are further apart than a specific cutoff distance given by their sizes (see the Supporting Information and Table S1 for details). In addition to the (truncated) repulsive Lennard-Jones potential between the crowders and the nucleotides, the potential energy of the system includes the backbone connectivity, hydrogen bonding, stacking, coaxial stacking, cross stacking, and excluded volume (see Supporting Information and Fig. S1 for details). The model treats electrostatic interactions with the Debye-Huckel potential (with effective backbone charges parameterized to reproduce the changes in duplex and hairpin stability as observed experimentally^34^).

For all the simulations performed in this work, we use the average-strength parameterization of the model, where hydrogen-bonding between A-T and C-G base pairs, as well as the stacking interactions, have interaction strengths independent of the base type and have been parameterized to reproduce the thermodynamics of DNA hairpins and duplexes with averaged AT and CG content^32,34^. Furthermore, to avoid any undesired base-pairing, we only allow hydrogen-bonding interaction between base pairs that are supposed to form in the duplex (or hairpin stems). The simulations are carried out in a cubic box of side length *L* = 17 nm with periodic boundary conditions, which corresponds to a strand concentration of 3.34 *×* 10^*−*4^ nm. The volume fraction *ϕ* occupied by the crowders is defined as *ϕ* = *N*_cr_*V*_cr_*/L*^3^, where *N*_cr_ is the number of crowders and 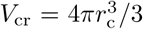 is the volume occupied by a spherical crowder of radius *r*_c_.

Before computing thermodynamics and kinetics quantities for duplex and hairpin formation, we wanted to check whether the diffusion of ssDNA in oxDNA in the presence of crowders was still Brownian. To do so, we first measured the mean-square displacement (MSD) of the center-of-mass of ssDNA of length 8 bases in crowded environments (Fig. S3) (*r*_c_ = 0.85 nm or 2.56 nm) with excluded volume fractions *ϕ* = 0.1 and 0.3, and observed diffusive behavior, with the MSD scaling linearly with time. The ssDNA diffusion is also not affected by the crowder mass (increasing the mass by a factor 27 does not change the measured diffusion coefficient of ssDNA). When we increase the excluded volume fraction from 0.1 to 0.3 we observe a decrease in the diffusion coefficient of ssDNA by approximately a factor of 1.6.

### B. Crowder sizes and volume fractions

In cellular environments, there are many types of molecules with various sizes. Large molecules, such as proteins with high molecular weight, will create macro-molecular crowding, while small molecules will have small-molecule crowding effects. To study the crowding effects across these scenarios, the excluded volume fraction of the crowding particles is selected from *ϕ* = 0.1 to 0.4 to mimic a cellular environment^17,18^. Additionally, the radius of the crowding particles is selected from 0.85 nm to 2.56 nm to mimic crowding conditions ranging from the small solutes to relatively large proteins.

### C. Simulation methods

Thermodynamic results in this work are obtained by using the virtual-move Monte Carlo (VMMC) algorithm^51^, which in combination with umbrella sampling (US) (see Supporting Information for details) allows efficient cluster Monte Carlo simulation of strongly-bound systems and was used previously to study the thermodynamics of DNA duplex and hairpin formations with oxDNA^32,35,49^. For the study of the DNA hybridization and DNA hairpin formation, we consider the strands (or stem) bound if there is at least one base pair present between the complementary native base pairs in the duplex (we do not allow for any mismatches). The melting temperature, *T*_*m*_, is taken to be the temperature at which the yield of duplex (or hairpin) is 50%. For the duplex system, we use its yield value extrapolated to the bulk^34,52^.

In order to study the kinetics of association, we use MD simulation with an Andersen-like thermostat^53^ with time-step 0.015 ps. To obtain good statistics for the association simulations, which are dominated by rare events, we used forward flux Nsampling (FFS). FFS facilitates sampling of a complex transition path ensemble by splitting a rare event into several intermediate stages. We first run MD simulations to estimate the flux, ϕ, through the first interface *λ*_0_. Simulations are then launched from states that have successfully crossed the first interface. The probability *P* (*λ*_1_ *λ*_0_) to reach interface *λ*_1_ from *λ*_0_ is estimated as the number of simulations that were started from a configuration at interface *λ*_1_ that successfully reached *λ*_2_ without crossing interface *λ*_*−*2_. Analogously, the probabilities are estimated for crossing the subsequent *N* interfaces *λ*_*i*_. The overall reaction rate constant *k* is estimated as 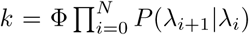. The definitions of respective interfaces for the study of duplex hybridization, hairpin formation, and strand displacement in crowded environments, along with a detailed description of the method, are provided in the Supporting Information and Fig. S2.

All MD simulations were performed at 37 *°*C and 1 m salt concentration. For a crowder of radius *r*_c_, we set the mass in the MD simulation to *m*_*r*_ = *m*_0_(*r*_*c*_*/r*_0_)^3^, where *m*_0_ corresponds to the mass of a single DNA nucleotide in the oxDNA model, and *r*_0_ = 0.85 nm is the radius of the smallest crowder sphere considered. Hence, we keep the density of the spheres representing the crowders constant and adjust their mass appropriately. We further set the diffusion constant of each crowder of radius *r*_c_ to *D*_*r*_ = *D*_0_*r*_0_*/r*_c_, where *D*_0_ = 0.24 nm^2^ps^*−*1^ is the diffusion coefficient of a single DNA nucleotide in the simulations. We set *D*_0_ to a value that corresponds to about 100 times greater than what was experimentally measured for a 14-mer DNA duplex in water. We intentionally choose the value of *D*_0_ to increase the diffusivity of the particles in the simulations and speed-up the sampling of the transitions (see Supporting Information for details).

For both thermodynamic and kinetic simulations, we only allow base-pair formation between the corresponding base-pairs in the hairpin stem (or base pairs in the 8-mer). Hence no alternative secondary structures or binding between bases different than the native base-pairs is allowed in the simulation.

## III. RESULTS AND DISCUSSION

### A. Crowding effects on DNA duplex and hairpin thermodynamics

We first study the crowding effects on the thermodynamic stability of an 8-mer DNA duplex and hairpins with a 6-nt stem and a 10-nt loop. The former reaction is inter-molecular while the latter is intra-molecular, covering the two most basic nucleic acid interaction types. We used VMMC simulations with oxDNA to obtain the melting temperature and free-energy profiles for both of these systems. The free energy between bound and unbound states, Δ*G*^0^, is computed as the difference between configurations with zero base pairs formed and those with at least 1 base pair formed, e.g., Δ*G*0 = *G≥*1 base pairs *− G*0 base pairs.

In the simulations, we considered different excluded volume fractions *ϕ* = 0.1, 0.2, 0.3, and 0.4. For each *ϕ*, we varied the crowder particle radius *r*_c_ from 0.85 nm to 2.56 nm with a step-size of 0.42 nm. The change in melting temperatures and free-energy profiles for the duplex and the hairpin as a function of number of base pairs formed are shown in Fig. 2. For the crowder with radius *r*_c_ = 0.43, nm, we only consider the volume fraction 0.1 (shown in Fig. S4A).

**FIG. 2.**
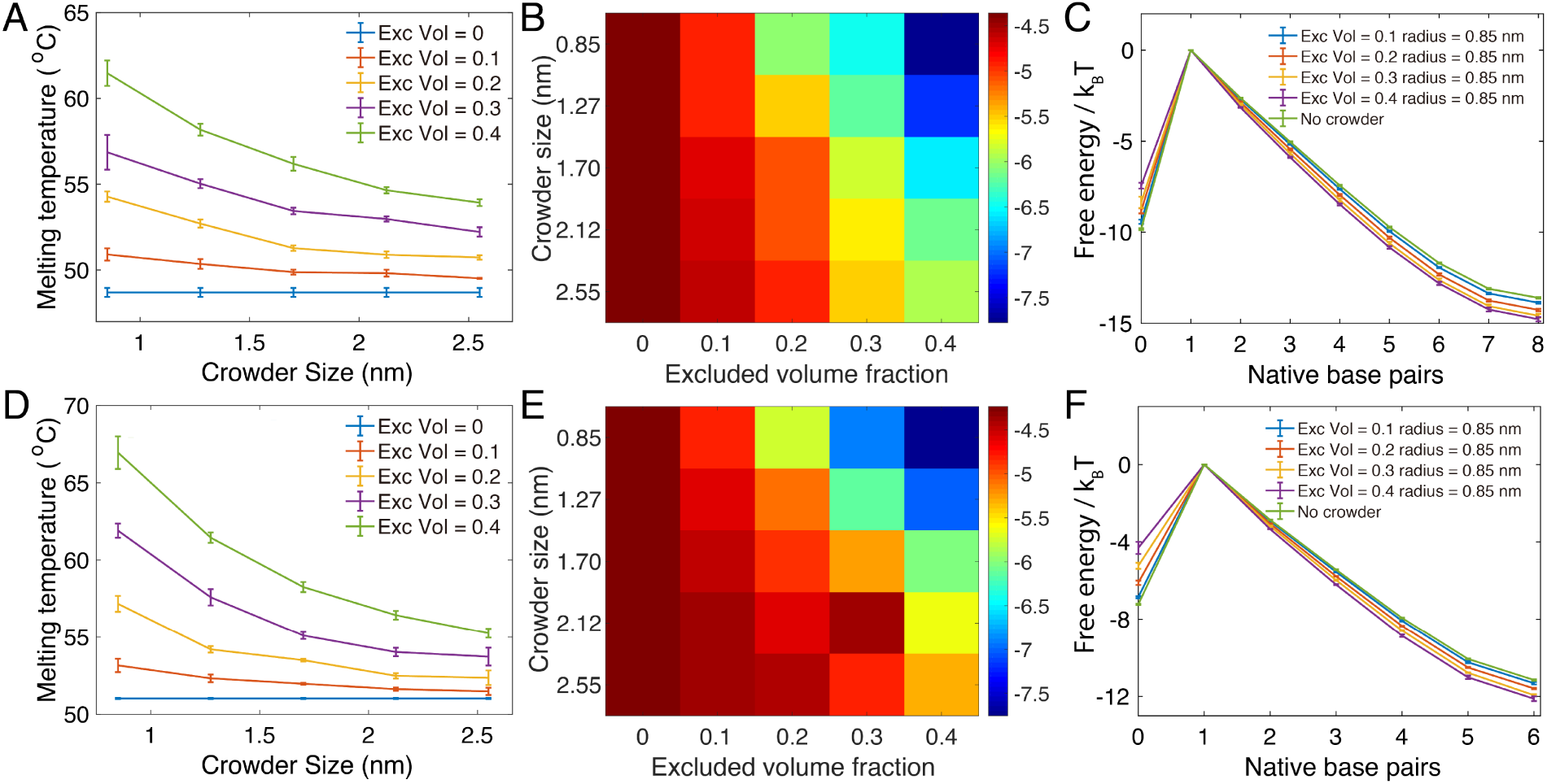
The thermodynamics of DNA duplex formation and hairpin closing in crowded environments. (A, D): The melting temperature of the 8-mer duplex (A) and hairpin (D) as a function of crowder radius *rc* under different crowding volume fractions *ϕ*. (B, E): The free-energy difference between the unbound and the fully-formed duplex (B) and the fully-formed hairpin stem (E). (C, F): The free-energy profile of duplex formation (C) and hairpin stem formation (F). In (A) and (D), the crowder-free case (blue) is shown alongside finite values of *ϕ* and *rc*.

Fig. 2A and D shows that the crowders enhance the stability of both duplex formation and hairpin closing, respectively, by increasing the melting temperature relative to the crowder-free environments. When the excluded volume remains constant, the melting temperature will decrease as the crowder radius increases. If the size of crowders remains the same, the larger the number of crowding particles present in the simulation box, the higher the the melting temperature, as the stability of the duplex or the hairpin is increased. The melting temperatures for the duplex and the hairpin without crowding particles are 48.70 *±* 0.26 and 51.02 *±* 0.04 *°*C, respectively. Under the largest crowder volume fraction and smallest crowder radius that we studied (*ϕ* = 0.4, *r* = 0.85), the melting temperatures increased to 61.47 *±* 0.74 and 66.95 *±* 1.05 *°*C, for the duplex and hairpin systems, respectively. Similar trends of the crowders’ effect are also seen in the free-energy changes of the duplex and hairpin formation. The heat maps in Figs. 2B and E indicate the free-energy difference between the single strand (0 bonds formed) and fully paired states (8 bonds for duplex and 6 for hairpin stem) under different *ϕ* and *r*_c_ for duplex and hairpin, respectively. For a fixed *ϕ*, the free-energy difference increases, favoring the fully formed duplex and hairpin, with smaller *r*_c_.

We further plot the free-energy landscapes versus the number of native base pairs formed for the 8-mer and for the hairpin in Figs. 2C and 2F, respectively, for *r*_c_ = 0.85 nm. In the plots, the free energy is set to zero when one base-pair is formed. The profiles for other studied combinations of *ϕ* and *r*_c_ are shown in the Supporting Information (Fig. S4 and S5). The landscapes all show that when the excluded volume fraction increases, the single-stranded state with zero base pairs formed becomes less favorable and at the same time the fully formed duplex (or hairpin stem) state is more stabilized (i.e. has lower free energy).

Fig. S4 and S5 also show the free-energy dependence on crowder radii *r*_c_ for the duplex and hairpin systems, respectively, varying from 0.85 nm to 2.56 nm at a fixed *ϕ*. We observe that the differences in the profiles between different *r*_c_ are larger with increasing excluded volume fraction. For a fixed volume fraction, the free-energy difference between states with zero base pairs formed and states with one base pair formed is smaller for smaller *r*_c_. Furthermore, the free-energy difference between a fully-formed duplex (8 base pairs) or a hairpin stem (6 base pairs) and the single base pair state is larger for smaller *r*_c_.

This behavior is in accordance with a theoretical analysis based on scaled particle theory (SPT)^48^, where the number density of the crowders enhances depletion effects through an increase in the effective osmotic pressure acting on the associating molecules. For fixed excluded volume fraction, the number density will be larger for smaller crowders, and thus their stabilizing effect is more pronounced.

The crowding-induced stabilization is further visualized by studying the equilibrium constant of duplex (hairpin) formation as a function of temperature for different crowding conditions. When the temperature varies, the reaction free-energy change can be deconstructed into its enthalpic and entropic contributions by the Van’t Hoff equations. Fig. 3 shows the plot of ln(*K*_eq_) vs 1/T for duplex formation, which is expected to have a linear relationship with a slope of Δ*H/k*_B_ and intercept of Δ*S/k*_B_. Fig. S6 shows the plots for hairpins and duplexes at a range of different crowder sizes. From the figures, we can see that the plots are all linear, as expected for crowders interacting only with excluded volume, with the straight line shifting to higher entropy as the excluded volume fraction increases for fixed *r*_c_, and shifting to lower entropy with an increase in *r*_c_ for fixed excluded volume fractions. The slope of the lines shows minimal or no change regardless of the change in *ϕ* and *r*_c_, as is expected for the stabilization of the duplex and hairpin systems coming from an entropic origin.

**FIG. 3.**
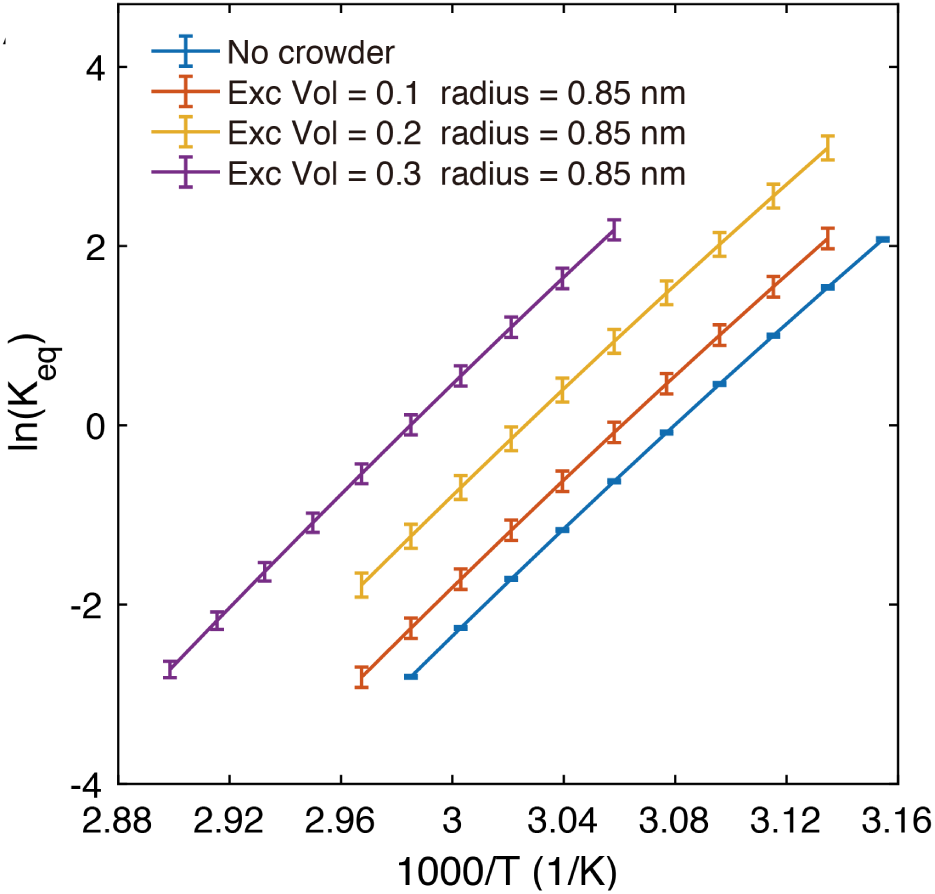
The Van’t Hoff plots for the 8-mer duplex. The entropic contribution is compared for a fixed *r*_c_ = 0.85 nm and *ϕ* ranging from 0.1 to 0.3.

### B. Comparing oxDNA thermodynamics with scaled particle theory

Scaled particle theory was developed to approximate the changes to the thermodynamics of polymer folding and assembly when crowded molecules are also present in solution at volume fraction *ϕ*^19,46,54^. It has previously been applied to both protein and RNA folding systems^28,55,56^. Here we apply this theory to DNA hairpin and duplex formation. In SPT the solution of DNA and crowder molecules is assumed to be ideal, and the ssDNA and the crowders are modeled as hard spheres having effective radii 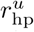 and *r*_*c*_, respectively^28^, as is illustrated in Fig. 4A(i) for the single strand. The folded hairpin is also modeled as a hard-sphere with radius 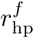 as is shown in Fig. 4A(ii). Fig. 4B(i) shows the 8-mer single strand in the duplex system with radius 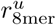, while in Fig. 4B(ii) the duplex is modeled as a hard spherocylinder with radius *r*_sc_ comprised of two adjacent, non-overlapping spheres that represent the two strands bound together. The radii of the unfolded single-strands are computed in oxDNA by measuring the end-to-end distance *R*_ee_ between terminal bases. See the Supporting Information for more details of the derivation. We compare the SPT models with our simulation results, as the coarse-grained crowder-oxDNA model provides a more detailed and physically accurate representation of double-stranded and single-stranded DNA, but is too complex to extract analytically the predicted free-energy changes as a function of the crowding environment.

**FIG. 4.**
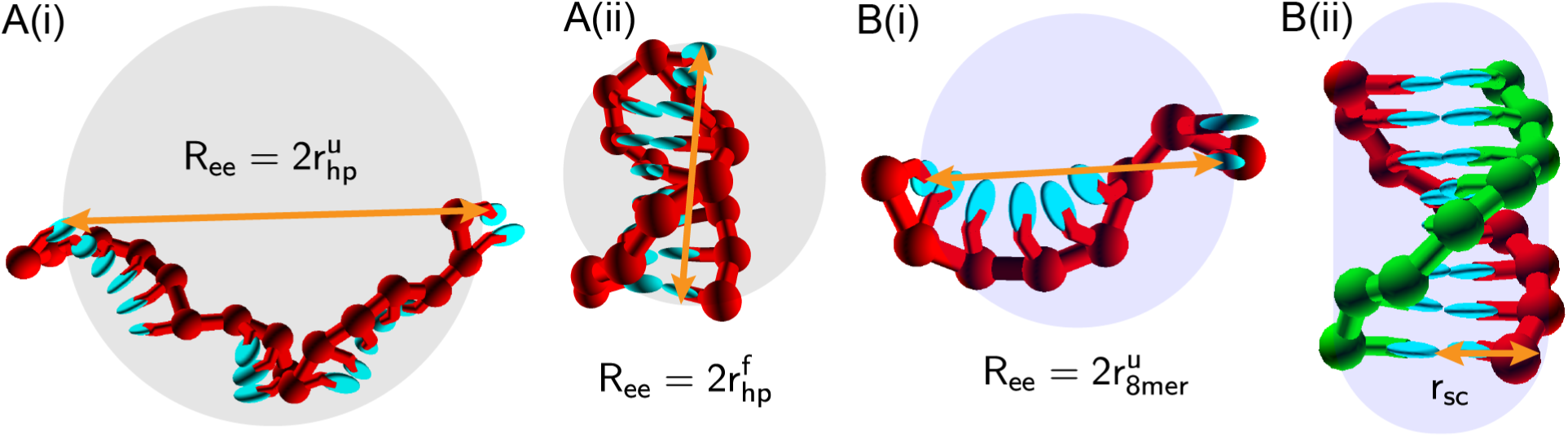
(a) The unfolded (i) and folded (ii) states of the hairpin system are illustrated, both showing the end-to-end separation *R*ee (orange arrow) and the radii of the hard sphere used to approximate the DNA in SPT. (b) The unfolded (i) and folded (ii) states of the 8-mer duplex system are shown. In (b)(i) only one of the complementary strands is shown with *R*ee (orange arrow) and the hard-sphere radii, while both strands are shown in the formed duplex in (ii) with *rsc* illustrating the spherocylinder radius (orange arrow). In all images spheres or the spherocylinder are not drawn to scale relative to the radii measured for the DNA. The superscripts ‘u’ and ‘f’ in the shown radii refer to “unfolded” and “folded” states, respectively.

In both duplex and hairpin systems, the quantity ΔΔ*G* is computed as the difference between the standard free-energies Δ*G*^0^ in crowded, and crowder-free environments

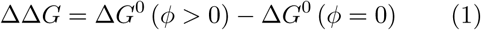

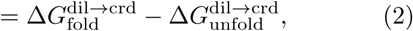

where Δ*G*^0^ is the free-energy difference between bound and unbound states in either system. Eq. 1 can be computed using SPT and with oxDNA (see the Supporting Information for details). For the hairpin system, expressions for both Δ*G*_fold_ and Δ*G*_unfold_ are derived in terms of *r*_hp_ for folded and unfolded configurations, respectively, the crowder radii *r*_*c*_, and the excluded volume fraction *ϕ*. An unfolded and a folded configuration are shown alongside each’s corresponding spherical representation (gray circles) in Fig. 5A(i) and A(ii) respectively. Similarly, ΔΔ*G*_duplex_, the difference in the free energy of duplex formation between environments with and without crowders, parameterized in terms of 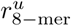 and *r*_sc_, can be calculated analytically for the duplex system under crowding conditions set by *r*_*c*_ and *ϕ*^56^. An unfolded single strand of length 8-nt is shown in Fig. 5B(i) with its spherical representation (blue circle), while in Fig. 5B(ii) a fully-formed 8-mer duplex configuration is shown with its corresponding spherocylinder representation.

**FIG. 5.**
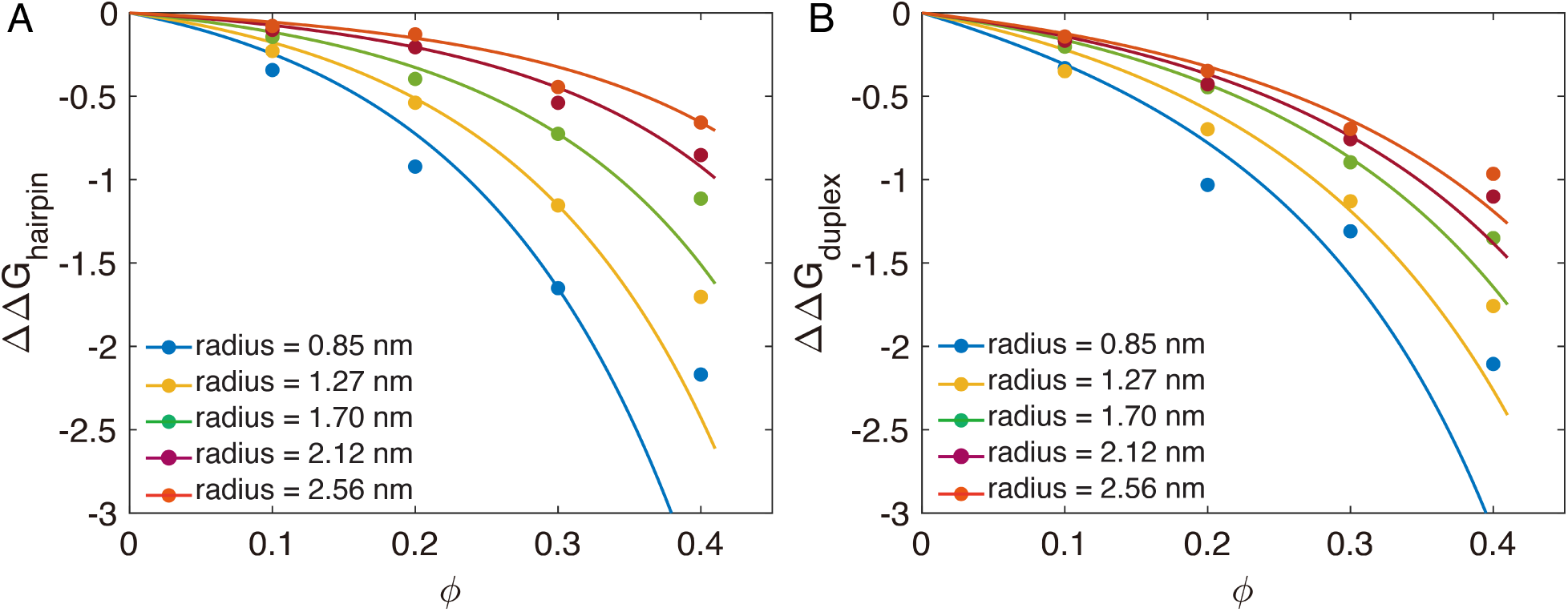
Scaled particle theory predictions (lines) are compared with the relative free energy ΔΔ*G* computed using oxDNA for (A) the 4-nucleotide loop hairpin (with 6 bp long stem) and (B) the 8-mer duplex (dots).

The radii of ssDNA and dsDNA abstracted as spheres and spherocylinders 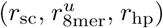 are used as free parameters to fit the SPT prediction to the measured ΔΔ*G* for hairpin and duplex in oxDNA simulations in environments with and without crowders, respectively (shown as dots in Fig. 5). To obtain the fits, Equations S17-S21 and S31-S32 were parameterized using these effective radius parameters, which are listed in Table S5.

Figures 5A and B compare the oxDNA computed ΔΔ*G*_*s*_ and the SPT predictions for the hairpin and the duplex systems with varying crowder sizes and *ϕ*. Even though our treatment of DNA in SPT is rather simplistic, in both systems, the SPT fit and oxDNA are in near-quantitative agreement for the larger crowder radii (*r*_*c*_ *⪆* 2 nm) and smaller crowder concentrations (*ϕ ⪅* 0.3). However, when *r*_*c*_ is less than 2 nm (smaller than the ssDNA), SPT predicts that the crowders influence the thermodynamics of the transition significantly more than the oxDNA model does. Additionally, when the crowder concentration is high, SPT predicts that the crowders influence the free energy of the transition more, as compared to oxDNA for both systems, though these differences are not as large when compared to the small *r*_*c*_ case. Hence, the SPT model provides as a good empirical model to predict the entropic effects of the crowders on duplex and hairpin stability but is not accurate enough for cases where the sizes of the crowders are comparable with the size of the individual nucleotide, where the approximation of the entire ssDNA chain as a sphere breaks down.

We further compare the fitted values of radii with 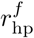 and 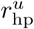 in the hairpin system, and 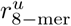 and *r*_sc_ with the values obtained from long sampling oxDNA MD simulations of hairpins and duplexes (Table S5). For the hairpin, the average strand end-to-end distance, *R*_ee_, was measured and used to compute *r*_*c*_ (Table S5). We found that the 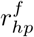 for the unfolded single strand in the hairpin system to be smaller than the 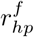 computed by oxDNA by about 25%. For the duplex system, the unbound single-strand radius measured by oxDNA was about comparable with the effective 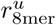, whereas the effective *r*_sc_ was smaller than the oxDNA computed quantity by about 25%.

We rationalize that the radii measurements obtained by oxDNA yield volumes for hard spheres that overestimate the true volume occupied by the DNA. This is because, in oxDNA, the ssDNA behaves as a freely jointed chain with excluded volume and does not have a fixed geometry. Similarly in the case of the duplex being modeled as a spherocylinder, DNA, in reality, is cylindrical with blunt ends and not capped-ends and so likely overestimates the true volume occupied by the duplex.

### C. Crowding effects on the kinetics of DNA hybridization

We next explore how the crowders affect the kinetic behavior of DNA during association using the crowder-oxDNA model. Specifically, we investigate the hybridization kinetics of the DNA 8-mer (Fig. 6), the hairpin formation with stem length of 6 and loop length of 10 (Fig. 7), and additionally a hairpin formation with stem length 6 and loop length 4.

**FIG. 6.**
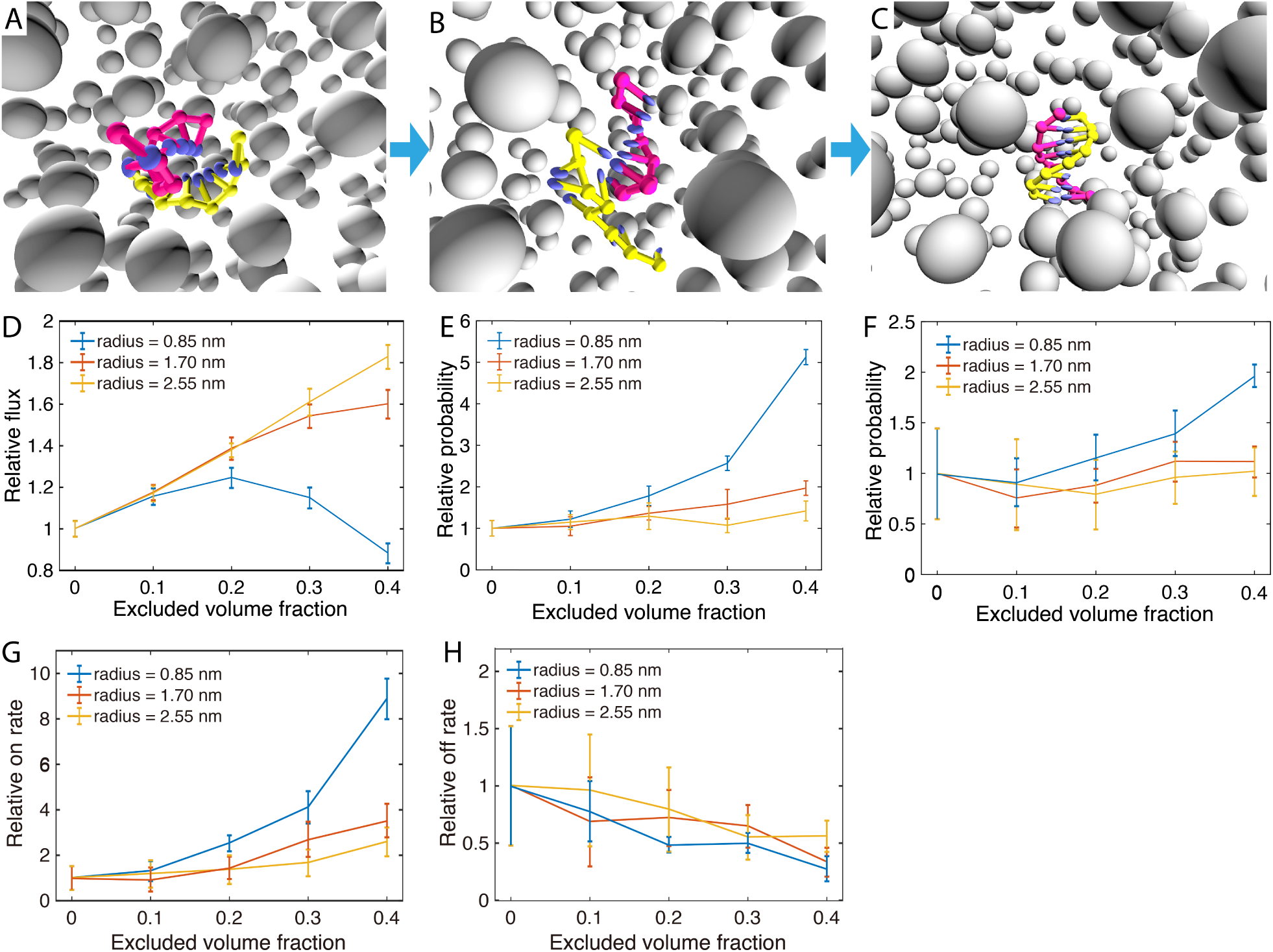
The kinetic FFS study of DNA 8-mer formation in different crowding conditions. (A-C) The illustrations show typical configuration at the three interfaces used to calculate transition rates for the hybridization process: (A) proximity between complementary bases, (B) the first base pair formed, (C) all 8 base pairs formed. The grey, isolated particles are crowders with the same radius, shown in perspective view. (D-F) The relative reaction flux/transition probabilities across the three interfaces, respectively. (G-H) The relative *k*_on_ and *k*_off_ for 8-mer duplex formation.

**FIG. 7.**
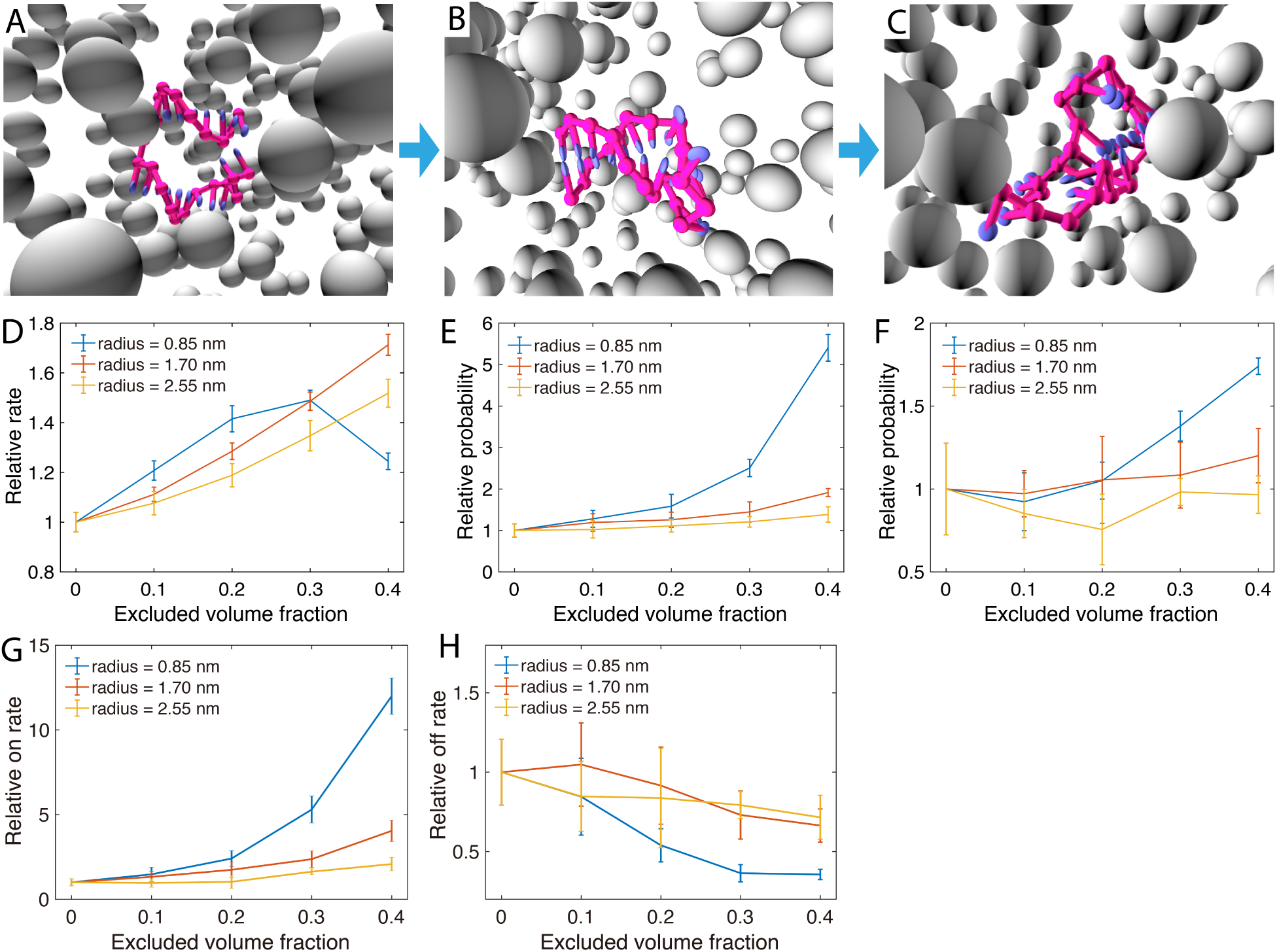
The kinetic study of hairpin closing with the crowder-oxDNA model. (A-C) A typical DNA hairpin formation process represented in three stages: (A) the stem bases getting close, (B) the formation of first base pair, and (C) the full base pairing of the stem. (D-F) The relative reaction rate of the three stages, respectively. (G-H) the overall relative *k*_on_ and *k*_off_.

We first simulate the duplex hybridization and hairpin stem formation with no crowders present using the oxDNA model. All reported rates, fluxes, and transition probabilities for crowded systems are normalized with respect to the mean values measured with no crowders present. Additionally, all kinetic simulations were carried out at 37 *°*C and with at a salt concentration of 1 µm.

We next perform kinetic studies of DNA in crowded environments with *r*_c_ = 0.85 nm, 1.7 nm and 2.56 nm. For each *r*_c_ considered, we study the kinetics of duplex and hairpin stem formation for *ϕ* = 0.1, 0.2, 0.3, and 0.4, respectively. Additionally, we only consider the hairpin with loop length 4 at *r*_c_ = 0.85 nm and *ϕ* = 0.1 and 0.4 (see Fig. S8). We use FFS, as described in the Methods section and in the Supporting Information, to sample the kinetics of duplex and stem formation, and obtain flux and transition probabilities.

The hybridization of DNA strands requires the diffusion of two complementary bases to spatial proximity, followed by the creation of a few base pairs to initiate a “zippering” up of the rest unpaired bases^26^. The 8-mer hybridization and hairpin closing processes are divided into three stages, schematically shown in Fig. 6A-C and Fig. 7A-C respectively. First, we calculate the flux through the first interface where the minimum distance between complementary bases is 0.85 nm. The relative normalized fluxes are shown in Figs. 6D and 7D, which compare different crowder conditions. We then sample the transition probabilities going from states where the minimum distance between complementary bases is *≤* 0.85 nm to those where one base pair has formed. The relative probabilities are shown in Figs. 6E and 7E for duplex and hairpin, respectively. The final interface, shown in Fig. 6F and 7F, measures the transition probability from one base pair formed to a fully-formed duplex or hairpin stem, respectively. The overall *k*_on_, obtained from multiplying the values obtained from the flux and transition probabilities, are shown in Fig. 6G and Fig. 7G for duplex and hairpin, respectively. For each case studied, we also obtained *k*_off_, which is shown in Fig. 6H and Fig. 7 for the two systems, by using the relation

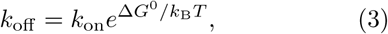

where Δ*G*^0^ was obtained as the free-energy difference between bound and unbound states (Fig. 2B and E) from VMMC sampling simulations under the same crowding conditions as in the FFS simulations where we measured *k*_on_.

Analysis of the binding kinetics shows similar trends both for the 8-mer duplex and the hairpin. Fig. 6D shows the relative fluxes (relative to that measured for the crowder-free case), which measure the rate at which complementary bases come into close spatial proximity. The plot shows that the flux increases linearly as a function of the excluded volume occupied by crowders with *r*_*c*_ = 2.56 nm. For *r*_*c*_ = 1.7 nm, the increase in flux is similar but becomes slightly slower when the excluded volume fraction *ϕ* reaches 0.3. However, for *r*_c_ = 0.85 nm, the flux shows non-monotonic behavior with increasing excluded volume fraction, first increasing until reaching a plateau at *ϕ* = 0.2, then decreasing thereafter. This phenomenon is likely due to competition between caging effects, which favor states at proximity, and increased viscosity caused by the crowders^20^. The confinement effect will help the DNA strands make contact more quickly since crowders fill up space in the system, and hence reduce the search space for complementary bases. At the same time, crowders also increase the viscosity and become obstructors that can hinder contact between complementary pairs. Similar effects are observed for the hairpin with loop length 10, where the flux for the smallest crowder (0.85 nm) decreases only after the volume fraction reaches 0.3 (Fig.7D). However, for the hairpin with loop length 4, the flux through the first interface decreases with increasing crowder volume fraction (Fig. S8), indicating that the obstruction effect dominates for hairpins with shorter loop lengths. We note, however, that the overall differences in flux between different volume fractions and crowder are all within a factor 2 for all cases studied.

After complementary base encounter each other over a small spatial separation, base-pairing may take place (Fig. 6A-B). For this transition, shown in Fig. 6-E, the relative transition probability increases for all crowder sizes considered with increasing crowder volume fraction. Since two DNA molecules are already spatially close to each other, the viscosity effect is likely not as important as the confinement effect on the nucleation of the hybridization.

At 40% volume fraction, the relative transition probabilities are 5.130. *±* 18, 1.970. *±* 17, and 1.42 *±* 0.24 for the three studied crowder radii, respectively. The reaction rate boost by the crowders mainly comes from the increased probability to create the first base pair. This supports the previously stated hypothesis^28^ that excluded volume effects lead to an increase in *k*_on_ by stabilizing the binding transition state.

Finally, in the last “zippering” step shown in Fig. 6F, the crowders have a slightly positive effect on the transition probability to the fully formed duplex, especially for the smaller crowders. The improvement may still come from the improved stability by the confinement effects.

The overall reaction on-rates, which are the product of the flux and respective transition probabilities of the three stages, are shown in Fig. 6G. For a fixed crowder radius, if we increase the excluded volume fraction *ϕ, k*_*on*_ will increase monotonically within the range of 0% - 40%. For a fixed *ϕ, k*_*on*_ will decrease with increasing crowder radius. Under the excluded volume of 40%, *k*_*on*_ for the crowder sizes considered are 8.87 *±* 0.89, 3.53 *±* 0.74, and 2.59 *±* 0.63, respectively.

We also calculated the off-rate *k*_off_ for comparison, which is shown in Fig. 6F. We plot the relative *k*_off_, normalized with respect to the value calculated for the system with no crowders. The relative off rates for different *r*_c_ are shown in Fig. 6H. The relative off rate decreases with increasing excluded volume fraction, but the crowding effects have smaller effects on *k*_off_ than on *k*_on_.

We further studied the effects of the mass of the crowder and the crowder diffusion coefficient on the association kinetics of the 8-mer, for crowders with radii *r*_c_ = 1.7 nm and excluded volume fraction *ϕ* = 0.2. As is shown in Fig. S7, we found that increasing the mass of crowders by a factor of 3, or increasing of diffusion coefficient by a factor of 3, has negligible effects on the measured association rate *k*_on_, and within error-bars, the mean *k*_on_ remains the same for all masses and diffusion coefficients considered.

The kinetics of hairpin formation is also thoroughly studied with the model, using the same FFS protocol as in the case of duplex formation. For the hairpin system we used the following interfaces for the flux and transition probability calculations: (i) complementary bases in the native stem region encounter each other over a small spatial range (Fig. 7A); (ii) the creation of one base pair in the stem region (Fig. 7B); and (iii) the fully-formed stem, which is when 6 base pairs have formed (Fig. 7C).

Overall, the crowder has a similar effect on hairpin closing that it had on duplex formation. The relative reaction rate *k*_on_ increases monotonously with an increasing excluded volume fraction. Under the highest excluded volume we simulated (40%), the relative reaction rates are 12.00 *±* 1.07, 4.04 *±* 0.63, and 2.08 *±* 0.40 for the crowder radii 0.85, 1.70, and 2.55 nm, respectively. Hence, the confinement effect has a slightly stronger influence on the intramolecular interactions (stem formation) compared to what was observed for inter-molecular duplex formation.

To understand how crowders influence hairpin folding when the loop length is varied, we also performed a kinetic study on a hairpin with a loop length of 4 nucleotides with *r*_c_ = 0.85 nm under different excluded volume fractions (0% - 40%), using the same FFS protocol that we employed for hairpin of loop length 10 (data are shown in Fig. S8). For the relative flux, the hairpin with a loop length of 4-nt shows a monotonous decrease with increasing excluded volume fraction. The probability of forming the first base pair from spatially proximal complementary bases in the stem is similar for excluded volume fractions 0.1 and 0.3. However, at *ϕ* = 0.4, the hairpin with loop length 10-nt has a higher relative reaction rate to form one base pair, while for the last interface (which measures the probability of stem formation given 1 formed base pair), the shorter loop has a slightly higher success probability than the 10-base loop. Overall, for crowder radius *r*_c_ = 0.85 nm, the *k*_on_ rate for hairpin formation is similar for the loop length 4 and loop length 10 at *ϕ* = 0.1 and *ϕ* = 0.2. However, *k*_on_ of stem formation with loop length 4 is smaller for larger excluded volume fractions (3.6 *±* 0.2 for *ϕ* = 0.3 and 6.5 *±* 0.4 for *ϕ* = 0.4) than *k*_on_ for the longer loop under the same crowding conditions. The results suggest that the crowders have stronger viscosity effects on the hairpin with the shorter loop length, as the first two stages of formation (reaching end-to-end proximity and forming the first base pair) are the same or lower for the short loop in all excluded volume fractions investigated. The short-loop hairpin also likely benefits less from the caging effects of the crowders.

We further use calculated Δ*G*^0^ for the hairpin with loop length 4 (Fig. S9) in combination with Eq. 3 to obtain the relative *k*_off_ at *r*_c_ = 0.85 nm. The quantity decreases by up to factor of 2.54 at 40% crowding fraction, similar to values obtained with the hairpin with loop length 10. Hence, the main difference in crowding effects between loop length 10 and 4 results from a change in *k*_on_.

### D. Strand displacement under crowded environments

Nucleic acid strand displacement is a reaction that allows one (invader) single-stranded DNA to displace another (incumbent) strand that is paired to a third (substrate) strand, as is illustrated in Fig. 8A-D. The invader strand (shown in red) will displace the incumbent strand (green) from the substrate strand (yellow) through a 6-nt toehold and then a branch migration process for 10-nt. Strand displacement processes are of great importance in active nucleic acid nanotechnology^6,57^ and are likely involved in several processes in biology^58^. To study how crowders affect the strand displacement reaction, we conducted both VMMC and FFS simulation for the strand displacement process to extract the thermodynamic and kinetic features of the reaction. In these simulations, the crowder radius was taken to be 1.75 nm at excluded volume fractions up to 40%.

**FIG. 8.**
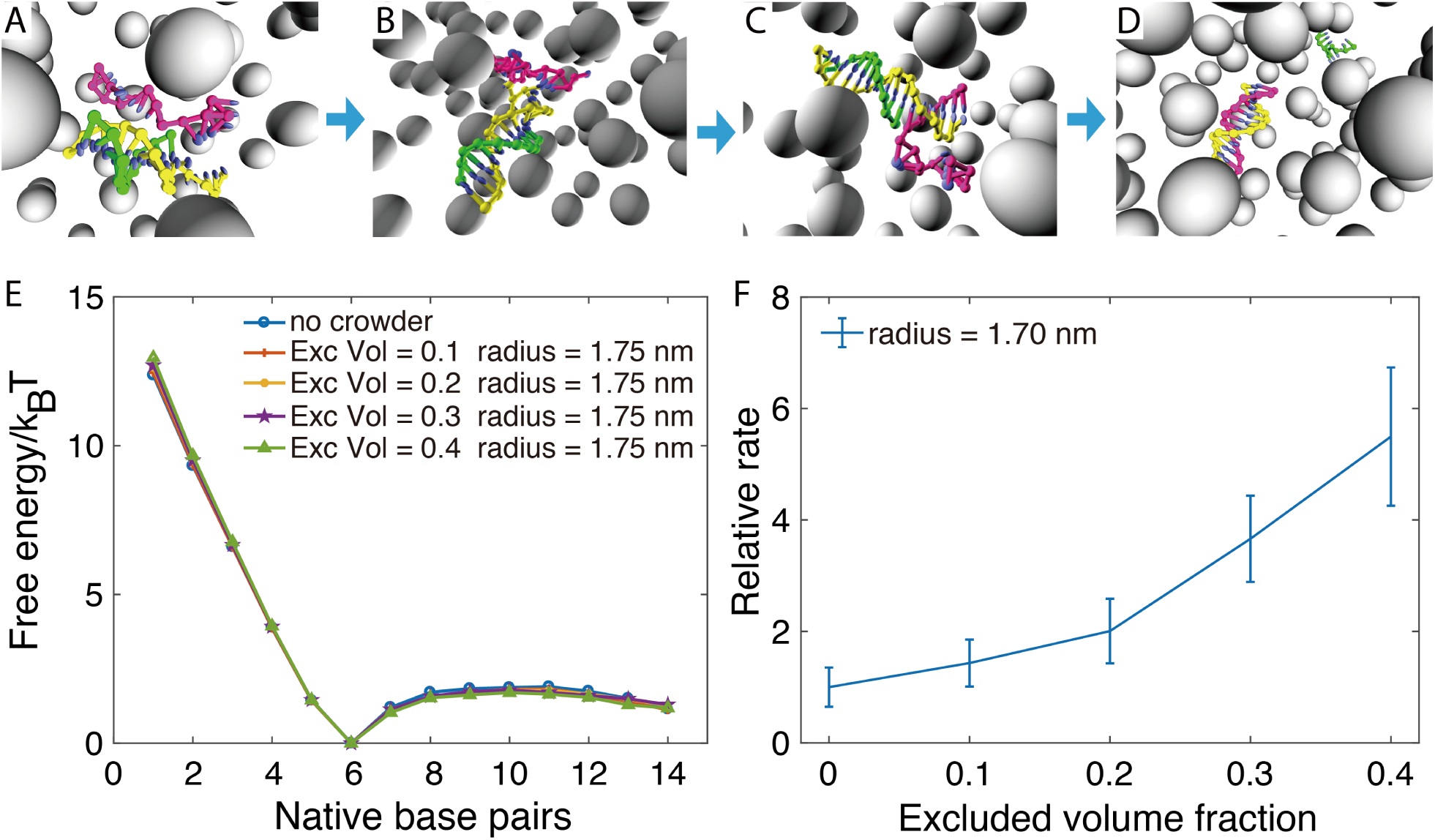
DNA strand displacement reaction in a crowded environment. (A-D) A snapshot of a typical configuration of the system crossing different interfaces: the contact of the toehold region, the first bond formation in the toehold region, the toehold region fully paired, and finally the full displacement of the incumbent strand. (E) The free-energy profile versus inter-strand base pair formation between the invader strand and the substrate strand. (F) The overall reaction rate of strand displacement under different excluded volume fraction with crowder radius of 1.7 nm

The free-energy profile of base pair formation between the invader strand and the substrate strand is shown in Fig. 8E, starting from the initial toe-hold association, to branch migration, and finally to the fourteenth base pair formed. Generally, the free-energy will first decrease after the first base pair forms as more stabilizing base pair contacts are created in the toehold association process, and then overcome a free-energy barrier to initiate the branch migration^44^. We also compared the free-energy profile in the toehold binding step with that of the 8-mer duplex hybridization, from 1 base pair to 6 base pairs having formed, and found it to be similar to the toehold binding profile (shown in Fig. S10). During the branch migration process, after crossing the initial barrier, the free-energy profile is mostly flat and does not show any significant change between the reaction under different excluded volumes.

To study how the crowders affect the kinetics of strand displacement, FFS is used to study the process by dividing it into four stages: (1) the contact of the toehold region, (2) the first base pair forming in the toehold region, (3) full base pairing in the toehold region, and (4) the branch migration process, as shown in Fig. 8A-D. The relative rate of the strand displacement as a function of excluded volume fraction is shown in Fig. 8F. The relative rate monotonously increases up to about 5.49 *±* 1.24 for *ϕ* = 0.4. The relative flux and transition probabilities for the individual interfaces are shown in Fig. S11, which show that the contribution to the overall augmentation in *k*_on_ with increasing *ϕ* comes from an increase in the binding rate of the toehold, the same as what was observed for the duplex formation kinetics with *r*_c_ = 1.7 nm. The presence of crowders has almost no effect on the branch migration kinetics (Fig. S11D) of the strand displacement reaction.

The overall study of the strand displacement in crowded environments suggests a possible strategy to increase the interaction rate of strand displacement cascades in DNA nanotechnology, by including crowders in the solution to speed-up the binding rate to the toehold region, thus accelerating the reaction.

## IV. CONCLUSION

In summary, in order to investigate DNA duplex hybridization and hairpin closing under crowded conditions, we have extended the previously developed coarse-grained oxDNA model by adding crowder interactions. The thermodynamic study revealed that crowders can stabilize DNA duplex and hairpin formation through purely an entropic origin. The kinetics simulations show that most of the change in free energy comes from speeding-up the binding rate, while the unbinding rate only changes slightly. These thermodynamic and kinetic trends are also consistent with prior experimental studies of RNA^28^, as well as with coarse-grained simulations for the folding of the WW peptide domain^59,60^. By dissecting the duplex and hairpin formation into a series of stages, we found that the reaction rate boost mainly comes from the stabilization of the intermediate transition state with one base pair having formed. However, we also note opposing effects, where the speed of association of complementary regions is partially hindered by collisions with crowders. Overall the stabilization of the transition state is the dominant effect and the association rate always increases with increasing occupied volume fraction. For a constant volume fraction, the effects of crowder sizes have also been explored, and we found that the thermal stabilization (and corresponding speed-up in *k*_on_) is larger for smaller sizes of crowders, in agreement with scaled particle theory (SPT) predictions.

Furthermore, we have provided a detailed comparison of the thermodynamics of hairpin and duplex formation with effective SPT that abstracts DNA strands and hairpins as as spheres and duplexes as spherocylinders. We find that in the regime of large crowders and small excluded volume fraction, SPT can fit simulation thermodynamics data with effective radius parameters, but the simplified spherical description assumed by SPT cannot reproduce behavior for excluded volume fraction over 30% and crowder sizes below 1.7 nm because it is no longer accurate enough to capture the entropic effects of the crowders on the DNA thermodynamics.

Nucleic acid interactions in cellular environments are complicated processes since the crowding particles have various shapes, sizes, and electrostatic properties. The coarse-grained model developed here can be further extended to include more interactions to mimic more complex systems, such as long-range electrostatic interactions, and be used to obtain insight into nucleic acid interactions in complex cellular environments. The present model, however, presents an important reference for comparison with other modeling and experimental efforts, as it captures the thermodynamic and kinetic effects solely based on excluded volume interactions with the crowders. Hence, any deviations from the simulation results have to originate from effects and processes not included in the current model, such as long-range electrostatic interactions or crowder-crowder and crowder-DNA binding affinities. Even though our analysis has been performed with a coarse-grained model of DNA, we expect the obtained results for duplex hybridization, hairpin formation, and strand-displacement would also apply to RNA molecules.

## V. ACKNOWLEDGEMENTS

We thank the members of the SŠulc group for helpful discussions and to E. Poppleton for help with the implementation of the visualization of the crowder-oxDNA model in the oxView tool. We thank Prof. J.M. Yuan for critical comments.

