## Supplementary Information for "Understanding DNA interactions in crowded environments with a coarse-grained model"

### Supporting Information for "Understanding DNA interactions in crowded environments with a coarse-grained model"

Fan Hong,[1,2] John S. Schreck,[1,3] and Petr Šulc[1,4,a)]
[1)] School of Molecular Sciences and Center for Molecular Design and Biomimetics at the Biodesign Institute, Arizona State University, Tempe, AZ 85287, USA
[2)] Wyss Institute, Harvard University, Boston, MA 02115, USA
[3)] Department of Chemistry, Drexel University, Philadelphia, PA 19104, USA
[4)] Center for Biological Physics, Arizona State University, Tempe, AZ 85287, USA

#### SI. Crower-oxDNA Model Details

OxDNA and its interaction potentials have been described in detail elsewhere[1–4]. In this work, we use the average-strength parameterization of the model from reference[4]. The model represents DNA as a 1D chain of nucleotides, where each nucleotide (sugar, phosphate and base group) is a rigid body with three interaction sites. The potential energy of the system can be decomposed as

$$V = \sum_{\langle ij \rangle} \left( V_{\mathrm{b.b.}} + V_{\mathrm{stack}} + V'_{\mathrm{exc}} \right) + \sum_{i,j \notin \langle ij \rangle} \left( V_{\mathrm{HB}} + V_{\mathrm{cr.st.}} + V_{\mathrm{exc}} + V_{\mathrm{cx.st.}} + V_{\mathrm{DH}} + V_{\mathrm{crow.\ exc.}} \right) \quad \text{(S1)}$$

where the first sum is taken over all nucleotides that are nearest neighbors on the same strand and the second sum comprises all remaining pairs. The interactions between nucleotides are schematically shown in Fig. S1. The backbone potential $V_{\mathrm{b.b.}}$ is an isotropic spring that imposes a finite maximum distance between backbone sites of neighbors, mimicking the covalent bonds along the strand. The hydrogen bonding ($V_{\mathrm{HB}}$), cross stacking ($V_{\mathrm{cr.st.}}$), coaxial stacking ($V_{\mathrm{cx.st.}}$) and stacking interactions ($V_{\mathrm{stack}}$) are anisotropic and explicitly depend on the relative orientations of the nucleotides as well as the distance between the relevant interaction sites. This orientational dependence captures the planarity of bases, and helps drive the formation of helical duplexes. The coaxial stacking term is designed to capture stacking interactions between bases that are not immediate neighbors along the backbone of a strand. Base and backbone sites also have excluded volume interactions $V_{\mathrm{exc}}$ and $V'_{\mathrm{exc}}$. Finally, we treat the electrostatic interactions using the Debye-Huckel approximation ($V_{\mathrm{DH}}$), with effective charges parameterized to reproduce the stability of short duplexes at sodium concentrations ranging from 0.1 to 1 M[4]. Hydrogen-bonding interactions are only possible between complementary (A-T and C-G) base pairs. In the average-strength parameterization that we use for all simulations, the strengths of interactions $V_{\mathrm{stack}}$ and $V_{\mathrm{HB}}$ are set to be the same for all types of nucleotides, parameterized to reproduce melting thermodynamics of short duplexes and hairpins with average-sequence content as predicted by SantaLucia's nearest-neighbor model[5].

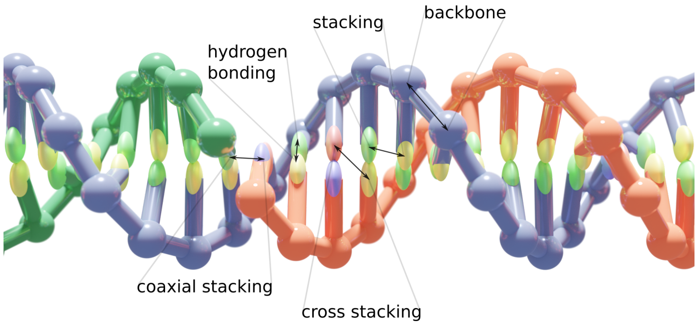


FIG. S1. **The oxDNA model**. A DNA duplex as modeled in oxDNA with labels corresponding to the coarse-grain potentials defining the force field.

To account for the presence of the crowders, we introduce a new interaction potential $V_{\mathrm{crow.\ exc.}}$ into the oxDNA model, that consists of the following terms:

$$V_{\mathrm{crow.\ exc.}} = V_{\mathrm{crowder\ crowder}} + V_{\mathrm{crowder\ back}} + V_{\mathrm{crowder\ base}} \quad \text{(S2)}$$

where the RHS of Eq. S2 correspond to the excluded volume interaction between two crowders, between a crowder and a nucleotide's backbone site, and between a crowder and the nucleotide's base site respectively. The functional form of the crowder excluded volume interaction potential is given by

$$f_{\mathrm{exc}}(r,\epsilon,\sigma,r^{\star}) = \begin{cases} V_{\mathrm{LJ}}(r,\epsilon,\sigma) & \text{if } r < r^{\star}, \\ \epsilon V_{\mathrm{smooth}}(r,b,r_{\mathrm{cut}}) & \text{if } r^{\star} < r < r_{\mathrm{cut}}, \\ 0 & \text{otherwise.} \end{cases} \quad \text{(S3)}$$

which consists of a Lennard-Jones potential function

$$V_{\mathrm{LJ}}(r,\epsilon,\sigma) = 4\epsilon \left[ \left(\frac{\sigma}{r}\right)^{12} - \left(\frac{\sigma}{r}\right)^{6} \right]. \quad \text{(S4)}$$

that is truncated using the quadratic smoothing function

$$V_{\mathrm{smooth}}(x,b,x^{c}) = b(x^{c}-x)^2, \quad \text{(S5)}$$

ensuring that the potential is a differentiable function that is equal to 0 after a specified cutoff distance $r_{\mathrm{cut}}$. We set $r^{*} = 2r_{\mathrm{c}}$ (where $r_{\mathrm{c}}$ is crowder radius) for the crowder-crowder interaction, and to $r^{*} = r_{\mathrm{c}} + r_{\mathrm{b}}$ for the interaction with the

---

[a)] Electronic mail:

**Crowder Interaction Parameters**

| | | | | | |
|---|---|---|---|---|---|
| Radius 0.85 nm | | | | | |
| $V_{\text{crowder}-\text{crowder}}$ | | | | | |
| $f_{\text{exc}}(\delta r_{\text{crowder}-\text{crowder}})$ | $\epsilon_{\text{exc}} = 2.00$ | $\sigma = 2.05$ | $r^\star = 2.00$ | $b = 113.8$ | $r_{\text{cut}} = 2.08$ |
| $V_{\text{crowder}-\text{back}}$ | | | | | |
| $f_{\text{exc}}(\delta r_{\text{crowder}-\text{back}})$ | $\epsilon_{\text{exc}} = 2.00$ | $\sigma = 1.4$ | $r^\star = 1.35$ | $b = 2897$ | $r_{\text{cut}} = 1.42$ |
| $V_{\text{crowder}-\text{base}}$ | | | | | |
| $f_{\text{exc}}(\delta r_{\text{crowder}-\text{base}})$ | $\epsilon_{\text{exc}} = 2.00$ | $\sigma = 1.22$ | $r^\star = 1.17$ | $b = 17111$ | $r_{\text{cut}} = 1.24$ |
| Radius 1.7 nm | | | | | |
| $V_{\text{crowder}-\text{crowder}}$ | | | | | |
| $f_{\text{exc}}(\delta r_{\text{crowder}-\text{crowder}})$ | $\epsilon_{\text{exc}} = 2.00$ | $\sigma = 4.05$ | $r^\star = 4.00$ | $b = 41.77$ | $r_{\text{cut}} = 4.09$ |
| $V_{\text{crowder}-\text{back}}$ | | | | | |
| $f_{\text{exc}}(\delta r_{\text{crowder}-\text{back}})$ | $\epsilon_{\text{exc}} = 2.00$ | $\sigma = 2.40$ | $r^\star = 2.35$ | $b = 88.50$ | $r_{\text{cut}} = 2.43$ |
| $V_{\text{crowder}-\text{base}}$ | | | | | |
| $f_{\text{exc}}(\delta r_{\text{crowder}-\text{base}})$ | $\epsilon_{\text{exc}} = 2.00$ | $\sigma = 2.22$ | $r^\star = 2.17$ | $b = 100.4$ | $r_{\text{cut}} = 2.25$ |
| Radius 2.56 nm | | | | | |
| $V_{\text{crowder}-\text{crowder}}$ | | | | | |
| $f_{\text{exc}}(\delta r_{\text{crowder}-\text{crowder}})$ | $\epsilon_{\text{exc}} = 2.00$ | $\sigma = 6.05$ | $r^\star = 6.0$ | $b = 25.0$ | $r_{\text{cut}} = 6.09$ |
| $V_{\text{crowder}-\text{back}}$ | | | | | |
| $f_{\text{exc}}(\delta r_{\text{crowder}-\text{back}})$ | $\epsilon_{\text{exc}} = 2.00$ | $\sigma = 3.40$ | $r^\star = 3.35$ | $b = 53.0$ | $r_{\text{cut}} = 3.44$ |
| $V_{\text{crowder}-\text{base}}$ | | | | | |
| $f_{\text{exc}}(\delta r_{\text{crowder}-\text{base}})$ | $\epsilon_{\text{exc}} = 2.00$ | $\sigma = 3.22$ | $r^\star = 3.17$ | $b = 57.4$ | $r_{\text{cut}} = 3.25$ |

TABLE S1. Parameter values in the model. All parameter values are in terms of the simulation units of energy and distance, with one length unit equivalent to 8.518 Å, and one energy unit to 41.42 pN nm.

backbone site or the base site, where $r_{\text{b}}$ corresponds to the radius of the backbone site or the base site in the oxDNA model respectively.

The parameters for each of the respective terms of $V_{\text{crow. exc.}}$ potential are listed in Table S1 for the crowder radii 0.85, 1.7 and 2.56 nm respectively. For interaction potentials shown in Table S1, $\delta r_{\text{crowder}-\text{back}}$ corresponds to the distance between the center of the crowder sphere and the backbone site on the nucleotide in the oxDNA model, and $\delta r_{\text{crowder}-\text{base}}$ is the distance between the center of the crowder sphere and the base site of the nucleotide. Finally, $\delta r_{\text{crowder}-\text{crowder}}$ is the distance between the centers of two crowder spheres.

As discussed in the main text, oxDNA has been extensively tested for other DNA properties and systems to which it was not fitted. Our success in describing all these phenomena gives us confidence to use it to study the dynamics of hybridization, hairpin formation, and strand displacement in the presence of crowders.

#### SII. Simulation Methods

##### A. Thermodynamics

###### 1. Virtual Move Monte Carlo

A standard approach for calculating the thermodynamic properties of computational models is the Metropolis algorithm[6]. A drawback with this approach is that only moving single particles at a time results in slow equilibration for systems with strong attractions. This is true for DNA strands, where collective diffusion is strongly suppressed if nucleotides are moved individually. Simulations can be made more efficient by using the Virtual Move Monte Carlo (VMMC), which allows for collective diffusion using cluster moves of particles[7]. Specifically, we use the variant presented in the appendix of reference 7. Initially, a particle is selected, and a move is chosen at random as in the Metropolis algorithm. The particle's neighbors are then added to a co-moving'cluster' with probabilities determined by the energy changes that would result from the move. Consequently, multiple particles tend to move at once. To use VMMC, we must select 'seed' moves of a single particle. For all VMMC simulations reported here, the seed moves were:

- Rotation of a nucleotide about its backbone site, with the axis chosen uniformly on the unit sphere and the angle drawn from a normal distribution with a mean of zero and a standard deviation of 0.22 radians.

- Translation of a nucleotide, where the displacement along each Cartesian axis is drawn from a normal distribution with a mean zero and a standard deviation of 0.15 simulation units of length (0.1277 nm).

##### 2. Umbrella Sampling

An important concept is that of a reaction coordinate (or order parameter) $Q$, which groups together microstates of a system that share some macroscopic property (for example, all configurations of strands with a certain number of base pairs). The free-energy profile as a function of $Q$ can provide useful information about the reaction, provided an appropriate choice has been made. Free-energy barriers can make certain regions of configuration space hard to reach, which prevents efficient sampling of all of the states of interest. The free-energy landscape can be artificially flattened by weighting states with different values of $Q$ appropriately, a technique known as umbrella sampling[8]. Thermodynamic properties of the system can then be extracted from simulations by unweighting the resulting distributions.

In particular, for an unweighted simulation a particular microstate with coordinates $\mathbf{q}^N$ and energy $E(\mathbf{q}^N)$ is sampled with probability

$$P(\mathbf{q}^N) \propto e^{-\beta E(\mathbf{q}^N)}. \tag{S6}$$

The equilibrium average of some variable $A(\mathbf{q}^N)$ is then given by the sum over all states, weighted by their Boltzmann factors:

$$\langle A \rangle = \frac{\int A(\mathbf{q}^N) e^{-\beta E(\mathbf{q}^N)} d\mathbf{q}^N}{\int e^{-\beta E(\mathbf{q}^N)} d\mathbf{q}^N}. \tag{S7}$$

By applying a weighting $w = w(Q(\mathbf{q}^N))$ to each value of the order parameter, we change the sampling frequency to

$$P_w(\mathbf{q}^N) \propto w(Q(\mathbf{q}^N)) e^{-\beta E(\mathbf{q}^N)}. \tag{S8}$$

where the subscript $w$ indicates a property of the weighted system. So we can artificially ensure that our simulation samples all states equally by making $P_w$ constant for all microstates. Equilibrium thermodynamic properties are then obtained by unbiasing afterwards, as can be seen by rewriting Eq. (S7) as follows:

$$\langle A \rangle = \frac{\int \frac{A(\mathbf{q}^N)}{w(Q(\mathbf{q}^N))} w(Q(\mathbf{q}^N)) e^{-\beta E(\mathbf{q}^N)} d\mathbf{q}^N}{\int \frac{1}{w(Q(\mathbf{q}^N))} w(Q(\mathbf{q}^N)) e^{-\beta E(\mathbf{q}^N)} d\mathbf{q}^N} \tag{S9}$$

$$= \frac{\langle A/w \rangle_w}{\langle 1/w \rangle_w}. \tag{S10}$$

Throughout this article, it makes sense to use the number of base pairs in our definition of $Q$. This is the usual choice for studying hybridization processes.

#### B. Kinetics

##### 1. Molecular Dynamics

Kinetic simulations were performed using an Anderson-like thermostat, similar to the one described in appendix A of reference 9. The Newtonian equations of motion for the system are integrated by Verlet integration[10] with a discrete time-step $\delta t$, so that the positions, velocities, orientations, and angular velocities of the nucleotides are recalculated at each time-step. This alone would give the DNA strands constant energy and cause ballistic motion. In reality, DNA in a solvent is being bombarded by water particles and thus undergoes Brownian motion. To model Brownian motion, the velocity of each nucleotide is resampled with a probability $p_v = 0.02$ from a Maxwell-Boltzmann distribution at the temperature of the solvent every 103 time steps. The algorithm also resamples angular velocities with a different probability $p_\omega = 0.0067$. The solvent thus acts as a large heat bath at a fixed temperature, ensuring that the simulated system samples from the canonical ensemble. On time scales longer than $N_{Newt}\delta t/p_v$, where $\delta t$ is the integration time step, the dynamics is diffusive. We choose $\delta t = 1.52 \times 10^{-14}$s for all dynamics simulations in this study. In oxDNA this time step gives a diffusion constant $D_{sim}$ for a 14-mer duplex that is about 100 times higher than experimental measurements[11] of $D_{exp} = 1.19 \times 10^{-10}\,\mathrm{m}^2\,\mathrm{s}^{-1}$.

This artificial increase in $D_{sim}$ is a common procedure for coarse-grained models where higher diffusion constants can be used to accelerate diffusion. Accelerated diffusion can also speed up certain processes by smoothing out, on a microscopic scale, energy profiles[12]. This can be advantageous because it means simulations utilizing coarse-grained models can be used to study more complicated systems. In a previous study using oxDNA, the hybridization kinetics of a non-repetitive sequence was considered[13]. In that study, it was shown that using higher friction constants (smaller diffusion coefficients) in simulations utilizing Langevin dynamics at 300K slowed down hybridization, but did not otherwise qualitatively affect the results. In particular, the tendency for initial base pairs to melt away rather than lead to a full-duplex was found to be preserved. Our systems are similar to those studied in reference 13, possessing similar numbers of total base pairing between the strands, and using a similar simulation temperature. Additionally, many approximations of real DNA have already been made in the construction of the oxDNA model, and we expect that running simulations with a diffusion coefficient that is larger than the experimentally measured value should preserve the effects that hairpins in single strands have on the relative hybridization and dissociation rates.

##### 2. Forward Flux Sampling

'Brute force' dynamics simulations using an Anderson-like thermostat are not efficient enough to generate a representative ensemble of trajectories that start in the single-stranded state and end in the duplex state. Thus, we resorted to using Forward Flux Sampling (FFS) to more efficiently calculate fluxes between local free-energy minima as well as sample the transition pathways. The term 'flux' from (meta)stable state A to state B has the following definition:

*Given an infinitely long simulation in which many transitions are observed, the flux of trajectories from A to B is*

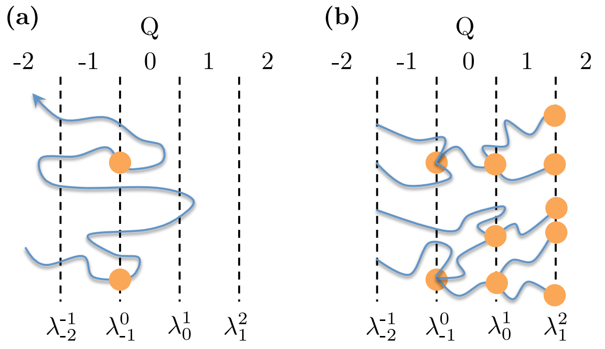


FIG. S2. (Color online) (a) Schematic illustration of the interfaces involved in flux generation. The flux is initially measured across the interface $\lambda^0_{-1}$. The orange dots indicate that a crossing by a trajectory contributes to the flux. These are also the states that are used to launch successive stages of the simulation. (b) In direct FFS, large numbers of configurations are randomly selected from the set that successfully crossed the interface $\lambda^0_{-1}$, and the probability of subsequently crossing the $\lambda^1_0$ interface (rather than returning to $Q = -2$) is measured. The process is then iterated over successively chosen interfaces until reaching $Q_{\rm max}$. Figure adapted from reference 13.

$\Phi_{AB} = N_{AB}/(\tau f_A)$, *where* $N_{AB}$ *is the number of times the simulation leaves A and then reaches B,* $\tau$ *is the total time simulated, and* $f_A$ *is the fraction of the total time simulated for which state A has been more recently visited than state B.*

FFS requires use of an order parameter, $Q$, which provides a descriptive measure of the extent of the reaction between states A and B. Additionally, the order parameter must be chosen such that non-intersecting interfaces $\lambda^Q_{Q-1}$ can be drawn between consecutive values of $Q$. At the beginning of an FFS implementation, a brute force simulation is run starting from states described by $Q$ = -2, and the flux of trajectories crossing the surface $\lambda^0_{-1}$ is measured. The total flux of trajectory from $Q$ = -2 to another free-energy minimum $Q = Q_{max}$ can be calculated as the flux of trajectories crossing $\lambda^0_{-1}$ multiplied by the probability that trajectories subsequently reach $Q = Q_{\rm max}$, all before returning to $Q$ = -2. The probability can be factorized as

$$P\left(\lambda^{Q_{\rm max}}_{Q_{\rm max}-1}|\lambda^0_{-1}\right) = \prod_{Q=1}^{Q_{\rm max}} P\left(\lambda^Q_{Q-1}|\lambda^{Q-1}_{Q-2}\right). \qquad \text{(S11)}$$

The first term in the product on the right-hand size of Eq. S11 is calculated by loading random configurations that have just crossed $\lambda^0_{-1}$, which are used to estimate $P(\lambda^1_0|\lambda^0_{-1})$. The process is then iterated for successive interfaces, and the flux as well as the trajectories that successfully reach $Q_{max}$ from the distribution of pathways can be measured.

#### SIII. Simulation Protocols

In this section, we discuss the implementation of the algorithms of Section SII for both single-stranded and duplex systems. We simulated the hairpin formation, duplex formation, and toehold-mediated strand displacement systems using FFS and VMMC simulations. We used a 'bonds' order parameter that measures the total number of base pairs, which can be specified to be intra- (for hairpins) or inter-strand (for duplexes and stand displacement) base pairs. The definition of a bonded base pair in our simulations is two bases with a hydrogen bonding energy below 0.596 kcal mol$^{-1}$. This value for the selected cutoff corresponds to about 15% of typical hydrogen-bond energy. In FFS simulations, we additionally used a 'distance order parameter' to measure the minimum distance between hydrogen-bonding sites overcorrect pairs of bases in the two strands.

##### 1. Duplex and Hairpin Thermodynamics

For the duplex and hairpin formation, we defined order parameters as the number of native bonds present in the stem for hairpins, and between two duplexes. The weights were manually adapted in order to assure proper sampling of all values of the order parameter. For each condition studied (as defined by crowder radius $r_{\rm c}$ and volume fraction $\phi$ occupied by crowders), we run at least three separate replicas of a free-energy calculation. The unbiased number of states for each value of the order parameter (using the procedure outlined in Sec. SII A 2) were used to plot the free-energy diagram. Histogram re-weighting was used to extrapolate free-energy profiles to other temperatures that the one at which the VMMC umbrella sampling simulations were run.

For each of the studied crowding conditions, we ran at least three independent umbrella sampling simulations. The error bars shown in the free-energy profiles in the main text in in the Supporting Information correspond to the standard deviation for the given value from all replicas for a given system. Each VMMC umbrella sampling simulation was run for at least $10^9$ steps.

For the melting temperature calculation in duplex formation simulations, care must be taken in extrapolating from a simulation of two strands to a bulk solution with many more strands, because fluctuations in local concentrations play an important role. If $\Phi$ is the ratio of bound to unbound states in a simulation of two molecules, the yield of a non-self-complementary duplex in a bulk solution (with the same average concentration of reactants) is given in reference 14 as

$$f^{\rm dim}_{\rm bulk} = \left(1 + \frac{1}{2\Phi}\right) - \sqrt{\left(1 + \frac{1}{2\Phi}\right)^2 - 1}. \qquad \text{(S12)}$$

The melting temperature occurs when $f^{\rm dim}_{\rm bulk} = 1/2$, which corresponds to a simulation yield of $\phi = 2$.

#### A. Kinetics

##### 1. FFS Simulation Details

In this section, we discuss the implementation of the FFS algorithm, discussed in Section SII. As mentioned in Sec-

| $Q$ | Description |
|---|---|
| -2 | $d > 3.40\,\text{nm}$ |
| -1 | $0.85\,\text{nm} < d \leq 3.40\,\text{nm}$ |
| 0 | $d \leq 0.85\,\text{nm}$ & $x = 0$ |
| 1 | $x \geq 1$ |
| 2 | $x \geq 8$ |

TABLE S2. The order parameter used in FFS simulations of duplex hybridization. The parameter $d$ is the minimum distance between any intended base pair on the two strands, $x$ is the number of base pairs between the two strands. A base pair is taken to be present if the hydrogen-bonding energy is less than $-0.596\,\text{kcal}\,\text{mol}^{-1}$ for x

| $Q$ | Description |
|---|---|
| -2 | $d > 3.40\,\text{nm}$ |
| -1 | $0.85\,\text{nm} < d \leq 3.40\,\text{nm}$ |
| 0 | $d \leq 0.85\,\text{nm}$ & $x = 0$ |
| 1 | $x \geq 1$ |
| 2 | $x \geq 6$ |

TABLE S3. The order parameter used in FFS simulations of hairpin closing. The parameter $d$ is the minimum distance between any intended base pair on the two strands, $x$ is the number of base pairs between the two strands. A base pair is taken to be present if the hydrogen-bonding energy is less than $-0.596\,\text{kcal}\,\text{mol}^{-1}$ for x

tion SII B 1, we simulated the duplex, hairpin, and strand displacement systems using molecular dynamics at $T = 37°$ °C. We also use the same definition of a bonded base pair that was discussed in Section SIII.

##### 2. Order Parameter Used in FFS Simulations

The order parameter used in the simulations are detailed in Tables S3, S2, and S4 for hairpins, duplex formation, and strand displacement respectively. Specifically, we use a combination of distance and base pairing criteria as outlined in Section SIII. Distance criteria are used to define states $Q = -2 \rightarrow 2$, and bonding criteria for states $Q =\geq 0$. In all the FFS simulations, the strands are designed to only allow for base pairs between correctly aligned native base pairs.

The flux is measured using a brute force simulations, counting the number of states per unit time that pass through interface $Q = 0$ coming from state $Q = -2$. The transition probabilities that we measure for hairpin (or duplex or strand displacement) correspond to transition through interfaces $\lambda_i^{i+1}$ for $i = 0$ and $i = 1$ (and also $i = 2$ for strand displacement).

| $Q$ | Description |
|---|---|
| -2 | $d > 3.40\,\text{nm}$ |
| -1 | $0.85\,\text{nm} < d \leq 3.40\,\text{nm}$ |
| 0 | $d \leq 0.85\,\text{nm}$ & $x = 0$ |
| 1 | $x \geq 1$ |
| 2 | $x \geq 6$ |
| 3 | $x \geq 16$ |

TABLE S4. The order parameter used in FFS simulations of strand displacement. The parameter $d$ is the minimum distance between any intended base pair on the two strands, $x$ is the number of base pairs between the two strands. A base pair is taken to be present if the hydrogen-bonding energy is less than $-0.596\,\text{kcal}\,\text{mol}^{-1}$ for x

#### SIV. Scaled Particle Theory

For reactions of the type P(solution) $\leftrightarrow$ P(aggregate), which include here the unimolecular hairpin folding and the bimolecular duplex formation reactions, the thermodynamic solubility constant absent crowders may be written[15]

$$\mathcal{K}_s^0 = \tfrac{1}{a_s} \tag{S13}$$
$$= \tfrac{1}{\gamma_s c_s} \tag{S14}$$

where $a_s$ is the activity, $\gamma_s$ is the activity coefficient, and $c_s$ is the concentration of ssDNA. When crowders are present ($\phi > 0$), the equilibrium constant $\mathcal{K}_s$ is taken to be proportional to that absent crowders as

$$\mathcal{K}_s \equiv \mathcal{K}_s^0 \gamma_s. \tag{S15}$$

In the main text we showed that both the thermodynamics and kinetics results results obtained from oxDNA simulations for a variety of crowder sizes and concentrations confirm that crowders do not significantly influence dissociation of hairpins or duplexes. As a result we may write

$$\ln\left(\frac{k_a}{k_a^0}\right) \approx \ln\left(\frac{\mathcal{K}_a}{\mathcal{K}_a^0}\right) = \ln \gamma_s \tag{S16}$$

where $k_a^0, k_a$ are the rate constants when crowders are absent and present at finite $\phi$, respectively.

##### A. Hairpin formation

In the case of the unimolecular reaction, an unstructured single-strand of DNA (said to be in state A) folds into a hairpin (said to be in state $A^*$). Both the crowder and the ssDNA are assumed to be hard spheres. For this reaction type, SPT may be used to compute the parameter $\gamma_1$ as[15]

$$\ln \gamma_1 = -\ln(1 - \phi) + A_1 z + A_2 z^2 + A_3 z^3 \tag{S17}$$

where

$$A_1 = R^3 + 3R^2 + 3R \tag{S18}$$
$$A_2 = 3R^3 + 4.5R^2 \tag{S19}$$
$$A_3 = 3R^3 \tag{S20}$$
$$z = \frac{\phi}{1-\phi} \tag{S21}$$

where $R \equiv r_i/r_c$, $r_i$ is the radius of a ssDNA ($i$ may be $hp$ for the strand that may fold into the hairpin, or 8-mer for either of the two strands that come together to form the duplex), and $r_c$ is the crowder radius.

#### B. Duplex formation

In the 8-mer system, two unstructured, complementary strands of DNA (taken to be states $A$ and $A^*$) come together to form a duplex (taken to be the state $AA^*$). The two unbound ssDNA in states $A$ or $A^*$ are again initially modeled as hard spheres having the same radii $r_s$. The duplex in state $AA^*$ is taken to be a hard spherocylinder comprised of two adjacent (hard-sphere) DNA strands with radius along the z-axis of the cylinder denoted $r_{sc}$, and the ratio of the axial length to the cylinder diameter is denoted as $L$. In reality, a ‘perfect’ DNA duplex, where strand $A$ is the reverse complement of $A^*$ and vice versa, is cylindrical in shape, but does not have capped ends, and is only ‘hard’ for lengths well below the persistence length of DNA (about 150 base pairs). The duplexes investigated here were only 8 bases long and as such were extremely rigid.

The activity of a spherocylinder may be estimated from Eqs. S17-S21 with[15]

$$A_1 = R^3 + 3R^2 + 3R + 1.5L(R^2 + R + 1) \tag{S22}$$
$$A_2 = 3R^3 + 4.5R^2 + 4.5L(R^2 + R) \tag{S23}$$
$$A_3 = 3R^3 + 4.5LR^2 \tag{S24}$$
$$z = \frac{\phi}{1-\phi}$$

and where $R \equiv r_{sc}/r_c$. The (oxDNA validated) assumption that only the forward rate is primarily influenced by crowders leads to the expression[15–17]

$$\ln\left(\mathcal{K}_a/\mathcal{K}_a^0\right) \approx \ln\gamma_1 + \ln\gamma_n - \ln\gamma_{n+1} \tag{S25}$$

where

$$\ln\gamma_{n+1} = \ln\gamma_r + \frac{d\ln\gamma}{dn} \tag{S26}$$
$$= \ln\gamma_n + \frac{d\ln\gamma}{dL}\frac{dL}{dn}. \tag{S27}$$

According to Eqs. S22-S24 we can write

$$\frac{d\ln\gamma_{sc}}{dL} = 1.5\left(R_{sc}^2 + 2R_{sc} + 1\right)z + 4.5\left(R_{sc}^2 + R_{sc}\right)z^2 + 4.5R_{sc}^2 z^3 \tag{S28}$$

where $R_{sc} = r_{sc}/r_1$. Assuming conservation of mass,

$$\frac{dL}{dn} = \frac{2}{3}\left(\frac{r_1}{r_{sc}}\right)^3 \tag{S29}$$

so that Eq. S25 may then be expressed as

$$\ln\left(\mathcal{K}_a/\mathcal{K}_a^0\right) = \ln\gamma_1 - \frac{d\ln\gamma_{sc}}{dL}\frac{dL}{dn} \tag{S30}$$

where the first term on the RHS is computed using Eqs. S17-S21, and the second using Eq. S28 and Eq. S29. Eqs. S25-S30 has been introduced for a general case where $n$ monomers can associate into a spherocylindrical shape, and in the case of a duplex, we only use them for $n = 2$.

For both hairpin or duplex formation reactions investigated here, the relative free-energy difference between crowder-free and crowded environments, $\Delta\Delta G$, relates to the equilibrium constants through

$$\Delta\Delta G_s \equiv \Delta G_s - \Delta G_s^0, \tag{S31}$$

which can be rewritten for the case of duplex formation as

$$\begin{aligned}\ln\left(\frac{\mathcal{K}_a}{\mathcal{K}_a^0}\right) &\equiv \Delta\Delta G_s \\ &= \Delta G_{AB}(\phi > 0) - \Delta G_{AB}(\phi = 0) \\ &- \Delta G_{A,B}(\phi > 0) + \Delta G_{A,B}(\phi = 0),\end{aligned} \tag{S32}$$

where the left hand side of Eq. S32 is computed according to SPT, while the right hand side is computed by estimating the free-energy terms using oxDNA simulations.

Following the approach taken in reference 18, we fit the SPT model to the oxDNA data by using the radius of the unfolded single-strand state in each hairpin and duplex systems as a fit parameter, and the spherocylinder radii $r_{sc}$ in the duplex system. The former measurement represents the “effective” radius of the unfolded states as the crowder size and volume fraction varies while the latter is the “effective” spherocylinder radius. We also ran long MD simulations to obtain representative trajectories from which we measured the end-to-end separation $R_{\text{ee}}$ in the single strand states (Fig. 4(a)(i) and (b)(i)), and the radius of the folded hairpin state (Fig. 4(a)(ii)) to compare the oxDNA predicted radii with the fitted effective radii. The sphero-cylindrical radius of the duplex, $r_{sc}$, is illustrated in Fig. 4(b)(ii).

For the unfolded single strands in Fig. 4(a)(i) and (b)(i), we measured the end-to-end distance $R_{\text{ee}}$ between terminal bases for the configurations in a trajectory to obtain an average measurement. For the hairpin shown in Fig. 4(a)(ii), we measured the average separation between a terminal stem base and a base in the middle of the loop, that is we measured the radius two times per configuration, then averaged them to obtain the effective radius of the folded structure.

In Table S5, we list the fitted radii used to compute the LHS of Eq. S32 and the computed radii from oxDNA simulations. The results of the thermodynamics comparison between SPT and oxDNA are shown in Figure 4 in the main text. As can be seen in Table S5, we observed no dependence of oxDNA-computed $R_{\text{ee}}$ on either the investigated values of $\phi$ or $r_c$ for both the hairpin and the duplex systems, in either folded or unfolded states. Additionally, the Table S5 also shows that as the effective radius of the unfolded single strands increases somewhat from smallest crowder to largest crowder, and is also independent of $\phi$ (not shown). Finally

| $r_c$ | $r^u_{hp}(*)$ | $r^u_{hp}$(oxDNA) | $r^f_{hp}$(oxDNA) | $r^u_{8-mer}(*)$ | $r^u_{8-mer}$(oxDNA) | $r_{sc}(*)$ |
|---|---|---|---|---|---|---|
| - | - | 2.08 ± 0.59 nm | 1.29 ± 0.13 nm | - | 1.22 ± 0.32 nm | - |
| 0.85 nm | 1.43 nm | 2.08 ± 0.59 nm | 1.29 ± 0.13 nm | 1.40 nm | 1.22 ± 0.32 nm | 0.86 nm |
| 1.28 nm | 1.55 nm | 2.08 ± 0.59 nm | 1.19 ± 0.13 nm | 1.23 nm | 1.23 ± 0.33 nm | 0.85 nm |
| 1.70 nm | 1.60 nm | 2.08 ± 0.59 nm | 1.19 ± 0.13 nm | 1.16 nm | 1.23 ± 0.33 nm | 0.85 nm |
| 2.13 nm | 1.60 nm | 2.08 ± 0.59 nm | 1.19 ± 0.14 nm | 1.14 nm | 1.23 ± 0.33 nm | 0.85 nm |
| 2.56 nm | 1.61 nm | 2.08 ± 0.59 nm | 1.29 ± 0.15 nm | 1.12 nm | 1.24 ± 0.33 nm | 0.85 nm |

TABLE S5. List of radius measurements used in the comparion of SPT with oxDNA thermodynamics. The parameter $r_c$ is the crowder radius, $r^u_{hp}(*)$ is the fitted radii of the unfolded hairpin state, $r^u_{hp}$(oxDNA) is the oxDNA measurement of the radius of the unfolded hairpin state, $r^f_{hp}$(oxDNA) is the oxDNA measurement of the radius of the folded hairpin state, $r^u_{8-mer}(*)$ is the fitted radii of the unfolded duplex state (referred to as 8-mer), $r^u_{8-mer}$(oxDNA) is the oxDNA measurement of the radius of the unfolded single strands that may come together to form the duplex, and $r_{sc}(*)$ is the effective radii of the spherocylinder. All oxDNA measured radii are mean values where the listed errors are the standard deviation in the mean.

the sphero-cylindrical radius of the duplex was observed to be nearly independent of crowder size or volume fraction, $r_{sc}(*)$ = 0.85 nm, significantly less than that measured previously with oxDNA where $r_{sc}$(oxDNA) = 1.15 nm.[1]

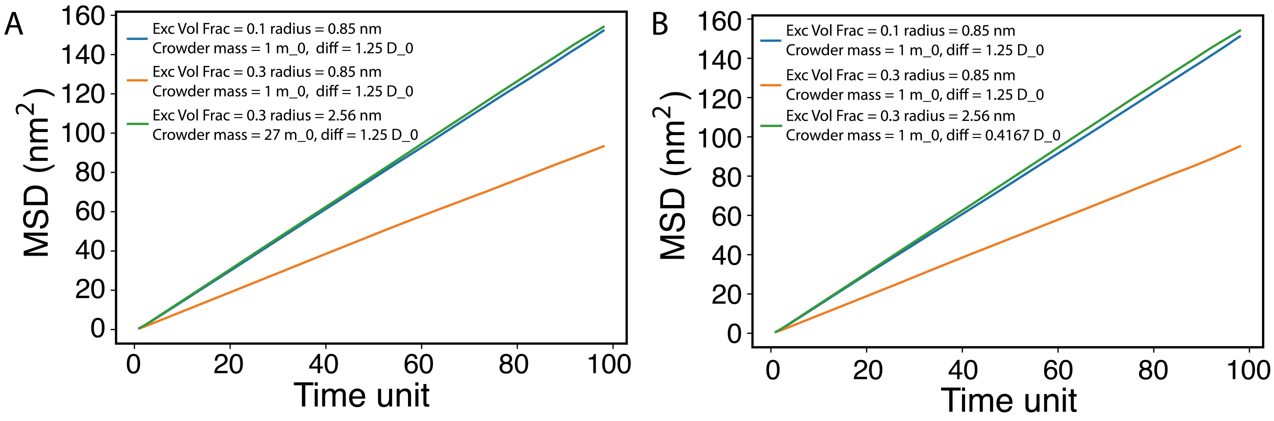


FIG. S3. The mean-square displacement (MSD) of the 8-mer ssDNA with crowders present. (A) Simulations with different masses of crowders. (B) Simulations with different diffusion coefficients of crowders. 1 time unit = 3.03 ps

.

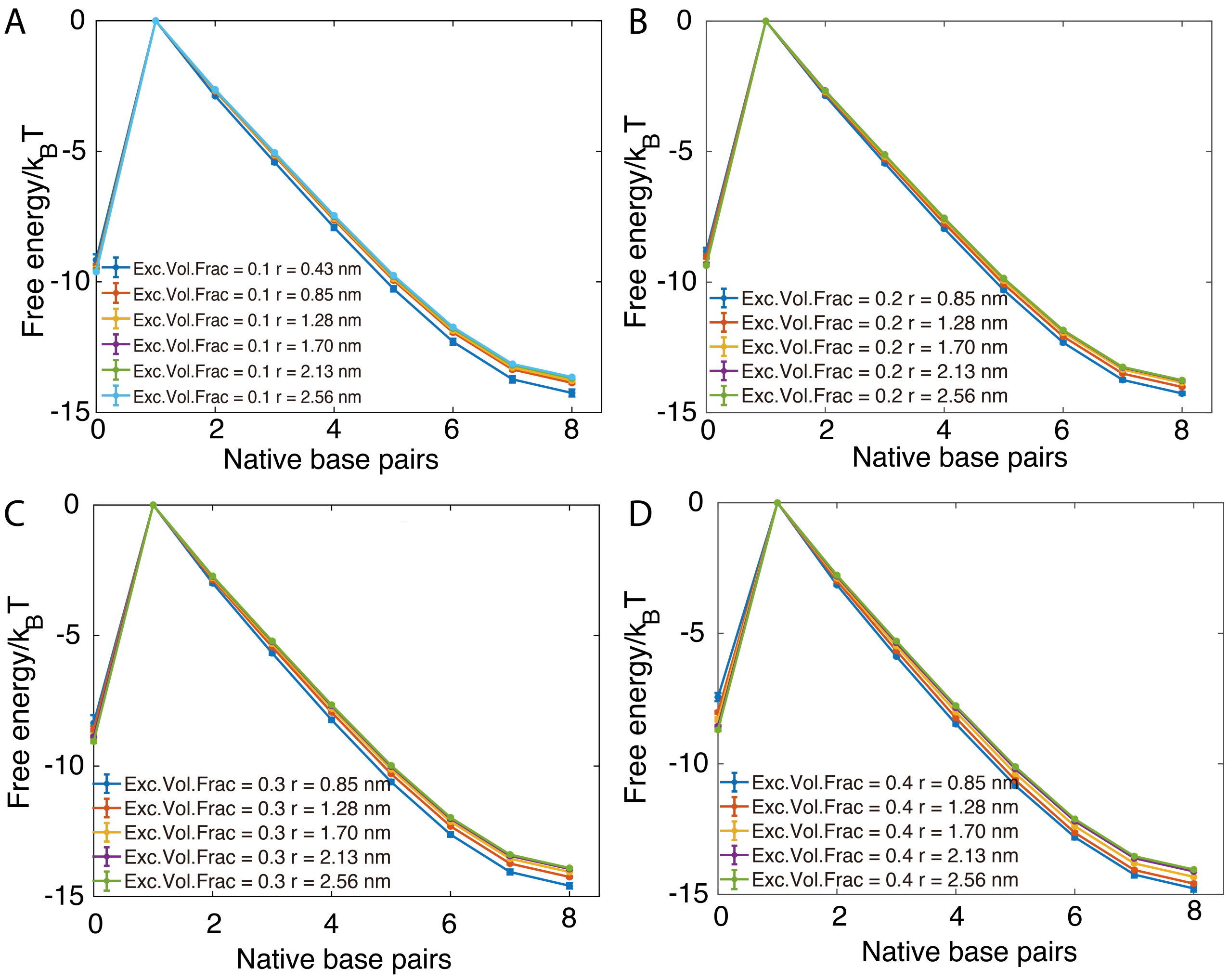


FIG. S4. The free-energy profile of the 8-mer duplex with excluded volume fractions of (A) 0.1, (B) 0.2, (C) 0.3, and (D) 0.4, respectively.

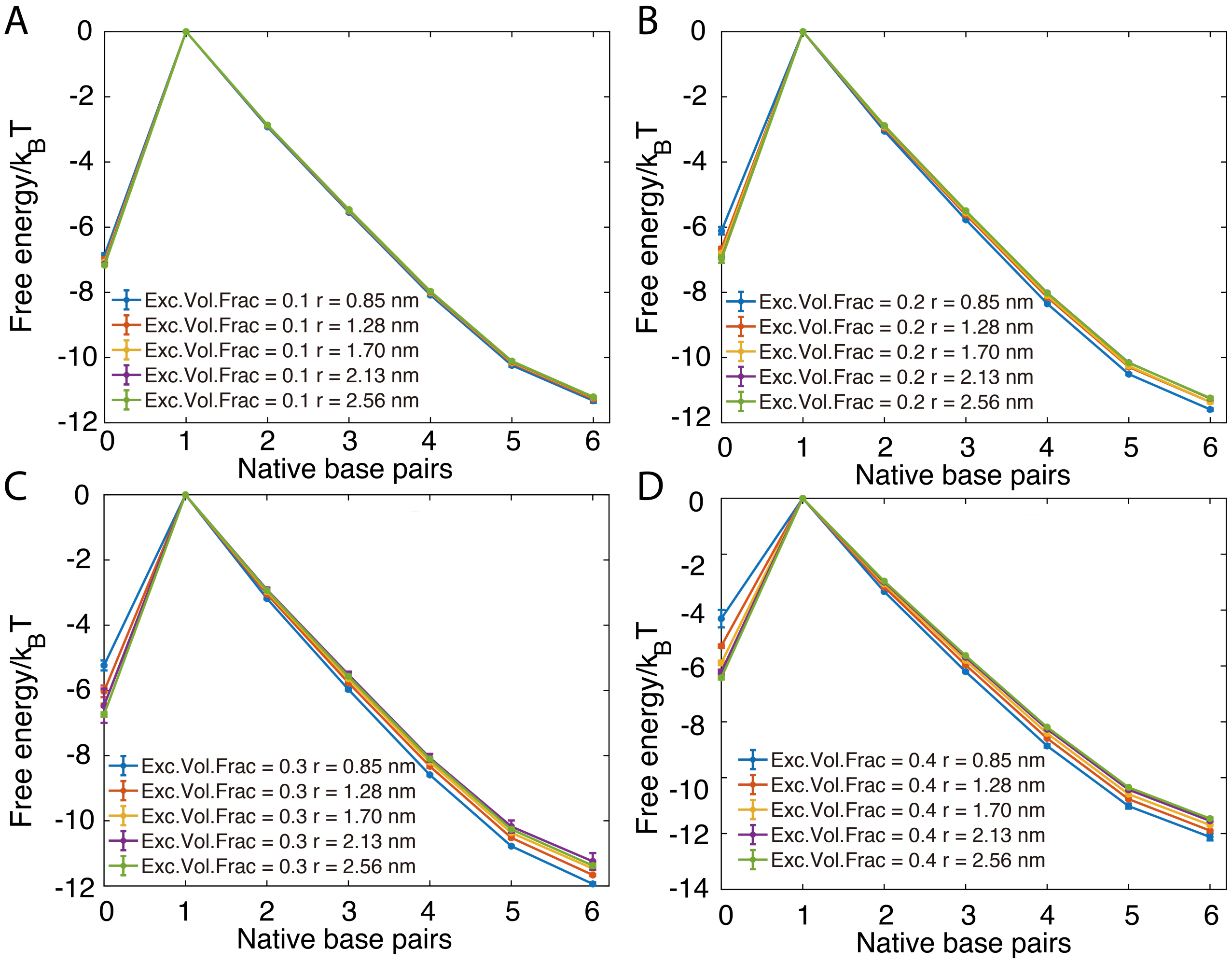


FIG. S5. The stem formation free-energy profile of a hairpin with a loop length of 10-nt. The panels show results where the excluded volume fraction was set to (A) 0.1, (B) 0.2, (C) 0.3, and (D) 0.4, respectively.

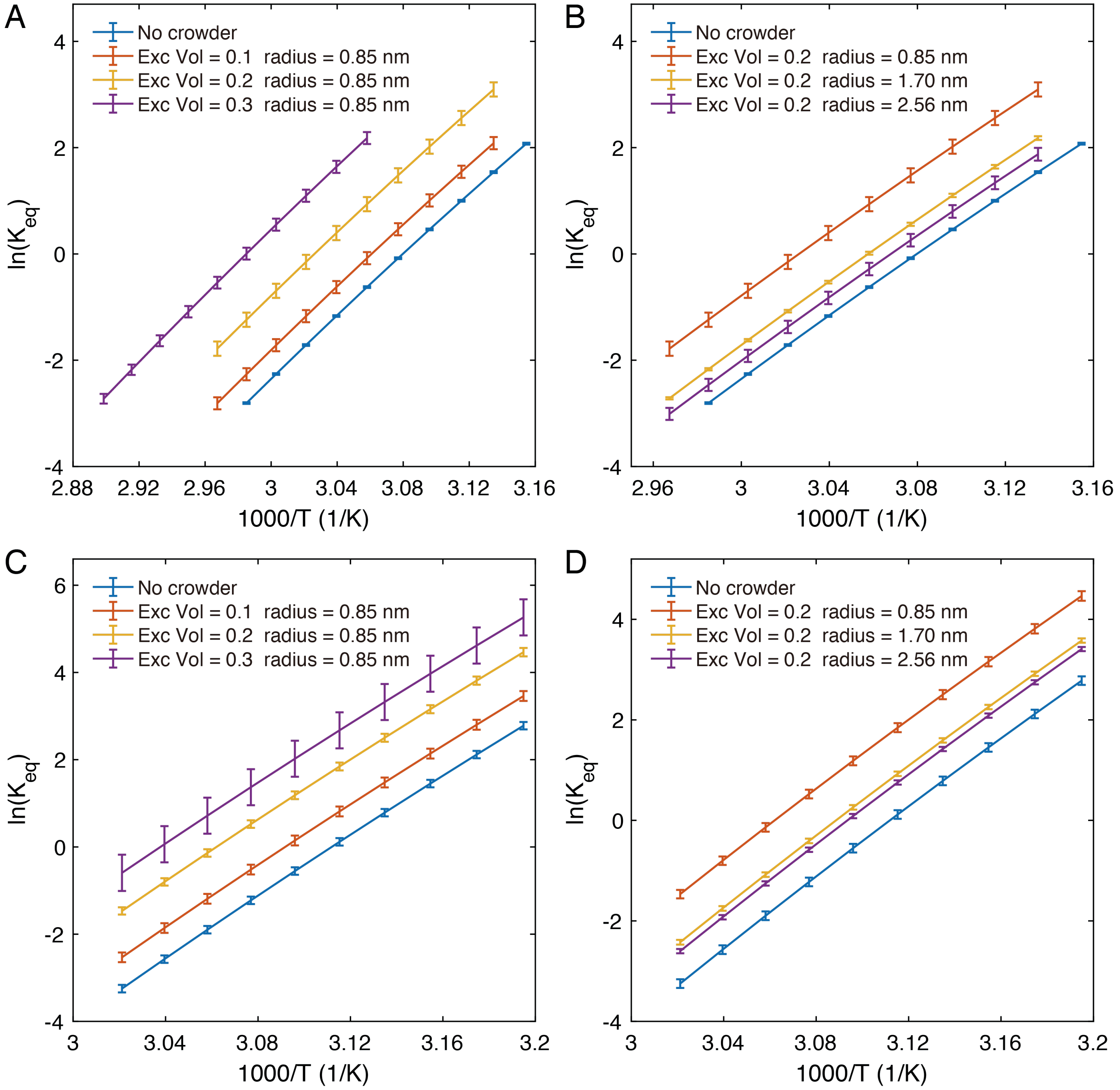


FIG. S6. The Van't Hoff plots of the DNA 8-mer formation (A and B) and hairpin formation with stem length 6 and loop length 10 (C and D). We compare the entropic contribution for a fixed crowder size of 0.85 nm with excluded volume fraction ranging from 0.1 to 0.3 (plots A and C), and for fixed excluded volume fraction 0.2 with radius ranging from 0.85 nm to 2.56 nm (plots B and D).

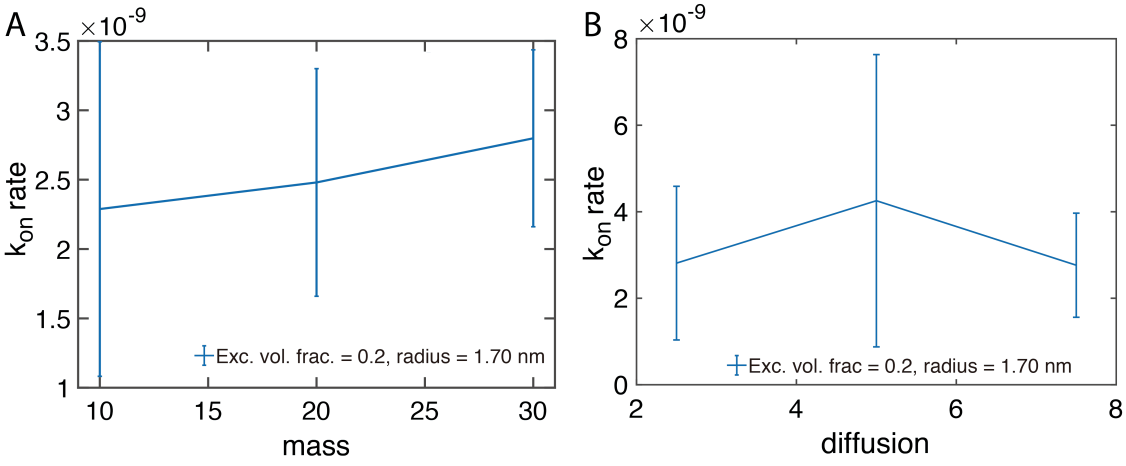


FIG. S7. The effects of crowder parameters on the DNA 8-mer formation. The excluded volume is 0.2 and the crowder radius is 1.70 nm. (A) The overall reaction rate on the change of crowder mass. (B) The overall reaction rate on the change of crowder diffusion. The change of mass and diffusion of crowding particles have very minor effects on the overall reaction rate of DNA hybridization.

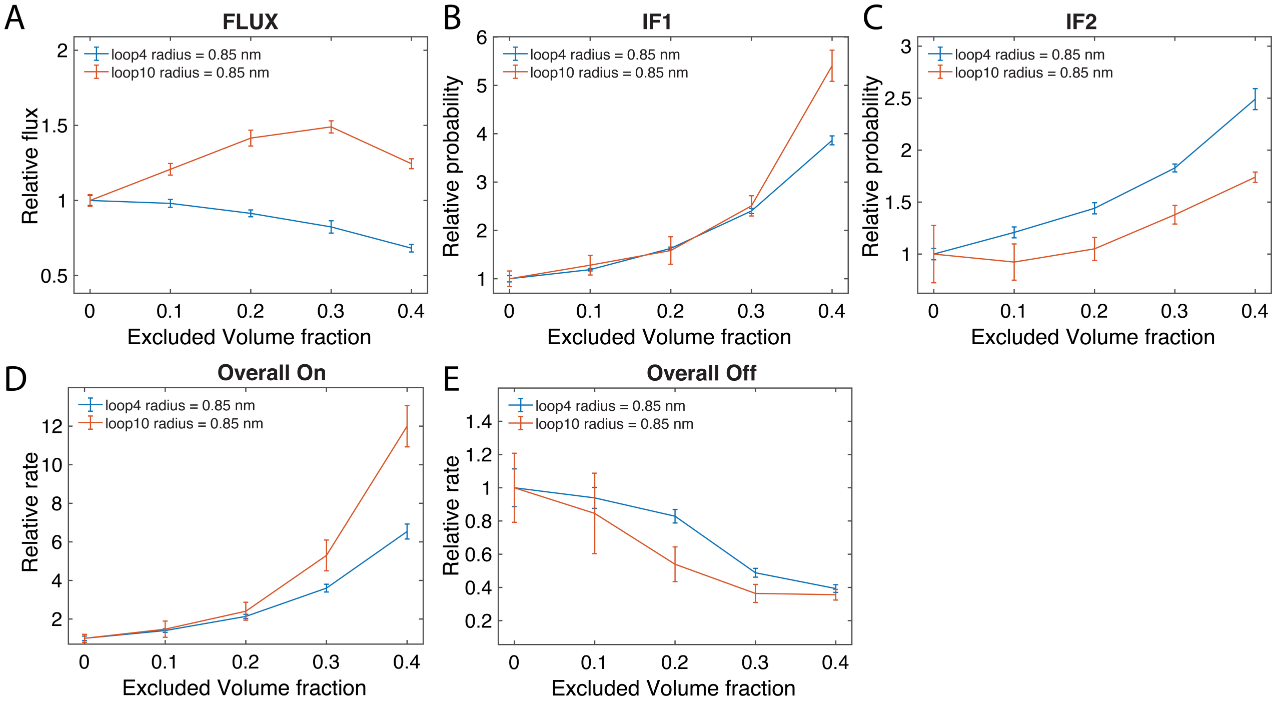


FIG. S8. The kinetic comparison between hairpin closing of loop lengths 4 and 6 for the flux (FLUX), IF1 (transition probability $P(\lambda_0^1|\lambda_{-1}^0)$) , and IF2 ($P(\lambda_1^2|\lambda_0^1)$) stages of the reaction. The overall relative $k_{\rm on}$ and $k_{\rm off}$ are also calculated for comparison.

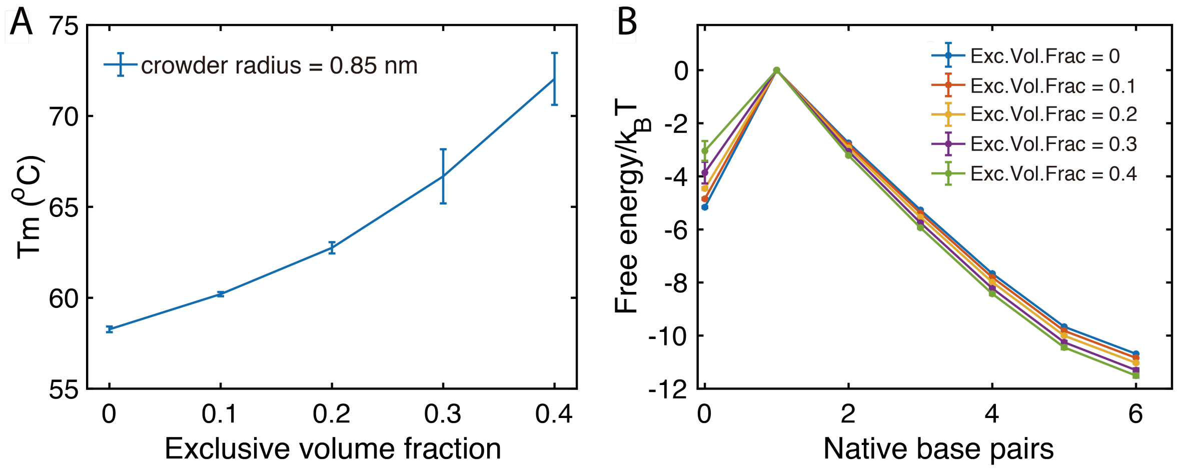


FIG. S9. The thermodynamics study of the hairpin with a loop length of 4-nt. (A) The melting temperature of the hairpin with a loop length of 4 versus excluded volume fraction. (B) The free-energy profile of hairpin closing versus native base pairs formed under various crowding conditions.

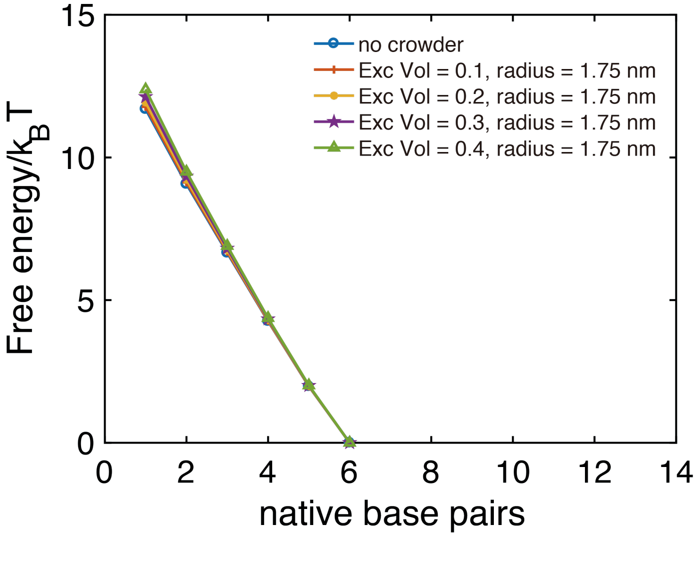


FIG. S10. The free-energy profile versus native base pairs formed for the DNA 8-mer hybridization, showing single base pair formation up to 6 base pairs having formed.

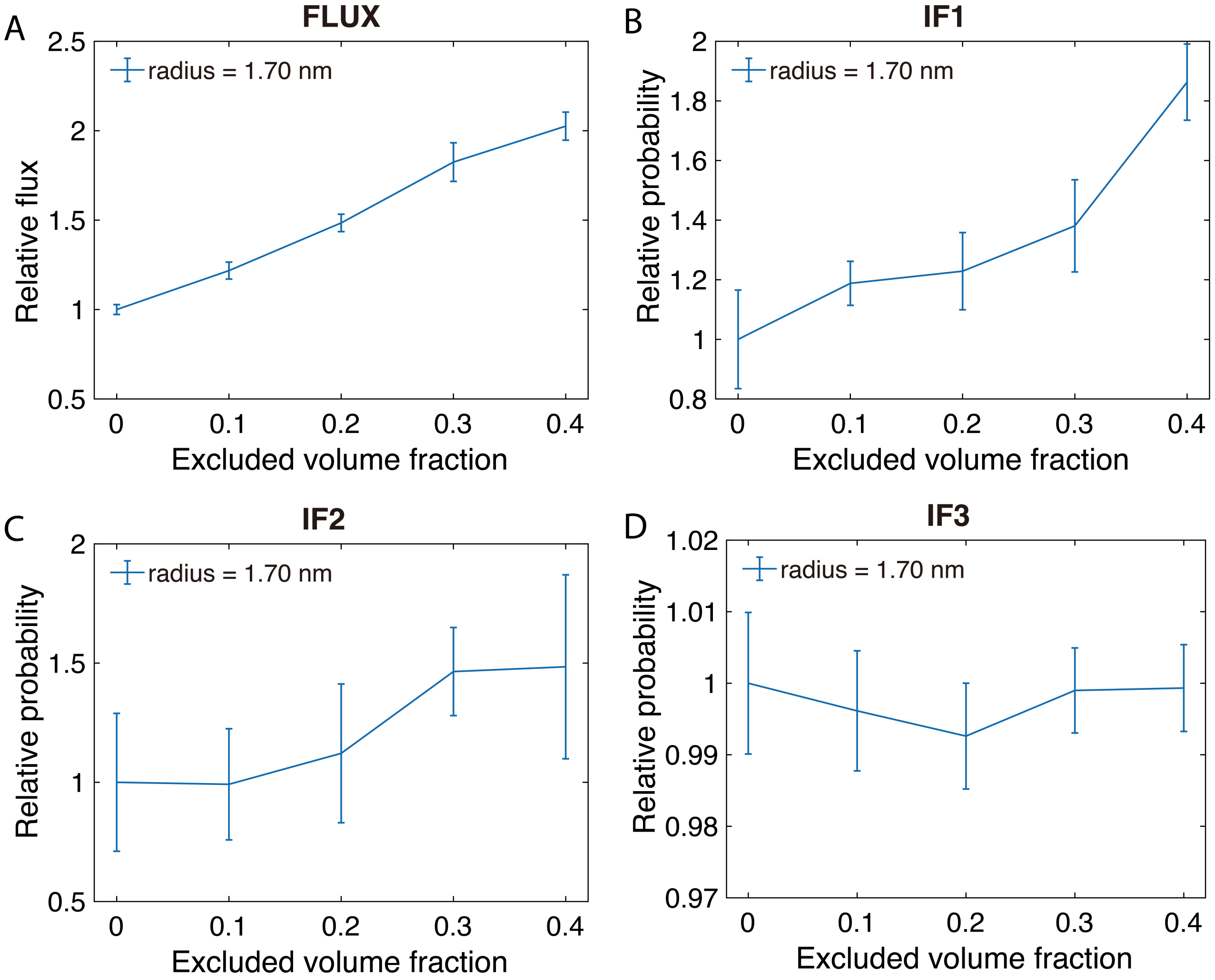


FIG. S11. The kinetic comparison of the strand displacement reaction at different excluded volume fractions. Showing flux (FLUX), IF1 ( corresponding to transition probability $P(\lambda_0^1|\lambda_{-1}^0)$), IF2 ($P(\lambda_1^2|\lambda_0^1)$), and IF3 ( $P(\lambda_2^3|\lambda_1^2)$)

#### SV. References


[1] Thomas E Ouldridge, Ard A Louis, and Jonathan PK Doye. Structural, mechanical, and thermodynamic properties of a coarse-grained dna model. *The Journal of chemical physics*, 134(8):02B627, 2011.

[2] T. E. Ouldridge. *Coarse-grained modelling of DNA and DNA nanotechnology*. PhD thesis, University of Oxford, 2011 [Published as a book by Springer, Heidelberg, 2012].

[3] Petr Šulc, Flavio Romano, Thomas E Ouldridge, Lorenzo Rovigatti, Jonathan PK Doye, and Ard A Louis. Sequence-dependent thermodynamics of a coarse-grained dna model. *The Journal of chemical physics*, 137(13):135101, 2012.

[4] Benedict EK Snodin, Ferdinando Randisi, Majid Mosayebi, Petr Šulc, John S Schreck, Flavio Romano, Thomas E Ouldridge, Roman Tsukanov, Eyal Nir, Ard A Louis, et al. Introducing improved structural properties and salt dependence into a coarse-grained model of dna. *The Journal of chemical physics*, 142(23):06B613_1, 2015.

[5] J. SantaLucia, Jr. A unified view of polymer, dumbbell, and oligonucleotide DNA nearest-neighbor thermodynamics. *Proc. Natl. Acad. Sci. U.S.A*, 17(95(4)):1460–5, 1998.

[6] D. Frenkel and B. Smit. *Understanding Molecular Simulation*. Academic Press Inc. London, 2001.
